# Hsa-miR-92a-3p regulates cell-cycle and signaling programs during human extra-embryonic lineage commitment

**DOI:** 10.64898/2026.04.29.721579

**Authors:** Pauliina Paloviita, Emilia Hautala, Rebecca Granskog, Sonja Nykänen, Roosa Pulkkanen, Ida Kirjanov, Reetta Santaniemi, Heli Grym, Christel Hydén-Granskog, Timo Tuuri, Sanna Vuoristo

## Abstract

MicroRNA (miRNA) levels increase during human embryonic genome activation (EGA) and have been implicated in the first lineage decisions. Re-analyzing single-cell and bulk small RNA-seq (sRNA-seq) from human and mouse embryos, together with new sRNA-seq of naïve human embryonic stem cells (hESCs) differentiated into hypoblast-like cells (HLCs) and extra-embryonic mesoderm (EXM)-like cells (EXMCs), we identify hsa-miR-92a-3p as highly expressed from oocyte to morula stage and dynamically regulated during HLC and EXMC formation. Functional inhibition of hsa-miR-92a-3p delays RACL (HLC/EXMC) differentiation, maintains epiblast-like and promotes trophectoderm (TE)-like transcriptional features, and reduces hypoblast and EXM marker acquisition. Transcriptome analyses revealed derepression of hsa-miR-92a-3p targets, including *FGF2*, and shifts in developmental and cell-cycle programs. The patterns of FGF protein stainings and flow cytometry-based cell-cycle analysis further implicate these pathways in RACL differentiation. Our findings position hsa-miR-92a-3p as a central regulator coordinating signaling and cell-cycle cues during extra embryonic lineage progression.

## INTRODUCTION

MicroRNAs (miRNAs) are ∼22-nt non-coding RNAs that post-transcriptionally calibrate gene expression, often acting on hundreds to thousands of mRNA targets to regulate developmental transitions and other cellular processes^1^. During the cleavage-to-morula stage, human embryos show increasing miRNA activity coincident with embryonic genome activation (EGA) and early lineage biasing^2–5^. Beyond small RNA transcriptional atlases^2,3^, recent surveys link intracellular miRNAs to inner cell mass (ICM) and trophectoderm (TE) lineage specification^3,6^, while secreted miRNAs have been implicated in implantation competence^7^. Together, these findings underscore the multifaceted roles of miRNAs in early human development.

miRNAs function by binding target transcripts, primarily through base pairing between nucleotides 2–7/8 of the miRNA, known as the seed region, and complementary sequences within the target mRNA^8^. Even noncanonical sites, such as 6mers or sites with seed mismatches, can induce weak repression when supported by strong 3′ pairing^1^, making target prediction based solely on sequence complementarity challenging. Although miRNA target sites are predominantly located within the 3′ untranslated regions (3′ UTRs) of mRNAs, functional sites have also been identified in 5′ UTRs and coding sequences^8^. miRNAs sharing the same seed sequence, seed family members, typically regulate overlapping sets of transcripts. Upon target recognition, miRNAs silence gene expression through cleavage and degradation or via deadenylation and translational repression^1^. Human miRNA loci are frequently organized into clusters that generate polycistronic transcripts containing multiple miRNA hairpins^9,10^. The chromosome 19 miRNA cluster (C19MC), the largest human miRNA cluster, is primate-specific, imprinted, and expressed almost exclusively in the placenta, with limited expression in human embryonic stem cells (hESCs) and certain tumors^11^. Recent studies show that C19MC members are enriched in human TE, yet their appearance as early as at the 8-cell stage (8C) and their broader expression across embryonic lineages suggest a role in supporting embryonic pluripotency^3^. Contrarily, the chromosome 14 miRNA cluster (C14MC), a conserved eutherian cluster, is expressed just before ICM–TE segregation and remains restricted to the ICM, underscoring the spatial and temporal precision of miRNA regulation in early human embryos^3^.

Because many miRNAs enriched in early human embryos are absent or divergent in rodents^12^, human pluripotent stem-cell (hPSC)–based models therefore offer a more accurate platform for studying miRNA-mediated lineage specification. Studies of naïve and primed murine and human ESCs have identified a core set of embryonic stem cell cycle (ESCC) miRNAs sharing a 5′-proximal AAGUGC motif^13,14^. These include members of the miR-371-373 and miR-302/367 clusters, which reinforce the ESC cell cycle, accelerate mesenchymal to epithelial transition during reprogramming, and counter TGFβ-induced epithelial to mesenchymal transition (EMT) through multi-target repression^13,15^. Members of the miR-17∼92 cluster, conserved across vertebrates and with the miR-106b/25 and miR-106a/363 paralogs in humans, also fall within this group^16,17^. Functionally, these miRNAs regulate cell cycle and proliferation, contribute to organ development, and, owing to frequent dysregulation in cancer, the miR-17∼92 cluster is widely known as ‘oncomiR-1’^16^.

Although 3D blastocyst-stage stem cell models (blastoids) have recently been profiled for miRNAs, analyses have primarily focused on TE- and ICM-like cells because hypoblast-like cells (HLCs), modeling human hypoblast cells that later form the yolk sac^18^, remain scarce in these systems^6,18^. Thus, while blastoids are valuable for modeling pre-implantation and implantation stages that are challenging to study with human embryos^19,20^, they are still suboptimal in recapitulating all the lineages. Recently developed stem-cell models now enable targeted generation of for instance HLCs and extra-embryonic mesoderm (EXM)-like cells (EXMCs)^21–24^, that model the EXM, which is involved in primary hematopoiesis, a source of extracellular matrix (ECM) proteins, and possibly a regulator of amnion and primitive streak^25,26^. This provides an opportunity to study miRNA programs in early lineages that remain uncharacterized in human development. Here, we systematically map miRNAs abundant in early human embryos and functionally test the role of hsa-miR-92a-3p in HLC and EXMC formation, demonstrating its critical role in the timely regulation of human embryonic and extra-embryonic specification in these rare cell types.

## RESULTS

### Hsa-miR-92a-3p and hsa-miR-372-3p are abundant in early human embryos

miRNA levels increased markedly from the 8C onward in human embryos^2,3^, preceding the first lineage segregation events, in which miRNAs have previously been implicated^3,6,13^. To define which miRNAs and seed families are most relevant during the first week of human development, we analyzed single-cell (sc) and embryo-level small RNA-sequencing (sRNA-seq) datasets and integrated these with functional assays in 2D and 3D differentiation models (**Fig. 1A**). We reanalysed previously published human sRNA seq datasets^2,27^ (**Fig. 1B-C, Supplementary Tables 1-4**) and noted that Yang dataset^27^ samples clustered primarily by developmental stage (**Extended Data Fig. 1A**). Yang oocytes and 2-cell stage (2C) embryos exhibited similar miRNA levels, whereas morulae showed a clear increase, consistent with miRNA upregulation during human EGA^2,3^ (**Fig. 1C**). Across these datasets, the seed families miR-302-3p/372-3p/373-3p/520-3p (fam-372) and miR-25-3p/32-5p/92-3p/363-3p/367-3p (fam-92) were among the most abundant from oocyte through morula stage (**Fig. 1B-C**). In both datasets, hsa-miR-92a-3p and hsa-miR-372-3p accounted for the majority of family-mapped reads, underscoring their prominence during the developmental window in which the first lineage decisions are established^4,5^.

**Figure 1.**
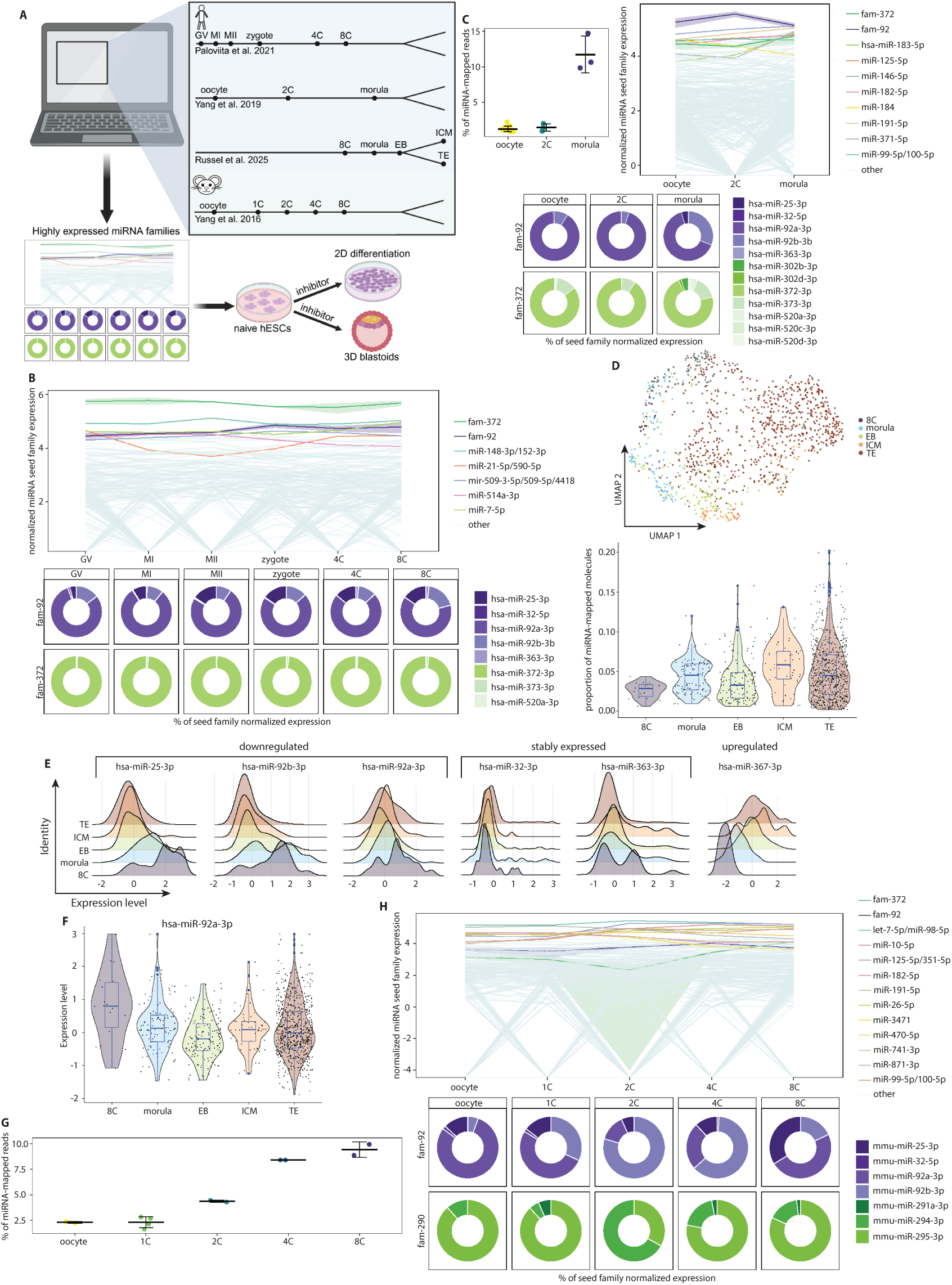
**(a)** Schematic overview of the study design. **(b-c)** Log-transformed mean TMM-normalized expression of miRNA seed families (colored) in human oocytes and early embryos ((b), Paloviita et al.^2^, Supplementary Tables 1-2; (c), Yang et al.^27^, Supplementary Tables 3-4), showing the five most highly expressed families at any stage (top) and the contribution (percentage of total family expression) of individual miRNAs within fam-92 and fam-372 (bottom). Ribbons for fam-92 and fam-372 indicate the standard deviation across developmental stages (top). For the Yang et al.^27^ dataset, the percentage of miRNA-mapped reads in each sRNA library is shown (c: top left), with dots indicating individual samples and vertical lines and whiskers indicating the group mean ± standard deviation **(d)** UMAP of Russell et al.^3^ single cell small-RNA-seq data with human embryo stages and lineages indicated by color (top) and the proportion of miRNA-mapped molecules per group shown as a blue boxplot overlaid on a violin plot (bottom). Boxplots show the median, 25^th^ and 75^th^ percentiles, and outliers; black dots indicate individual cells. **(e)** Ridge plots of miRNA molecule–proportion distributions (over total miRNA molecules) after data integration in the Russell et al.^3^ dataset for fam-92 members grouped as up-regulated, down-regulated, or stable between the 8-cell stage and Inner cell mass or trophectoderm. See Extended Data Figure 1c. **(f)** Boxplot (blue) overlaid on a violin plot showing normalized hsa-miR-92a-3p expression across human embryo stages and lineages^3^, colored by group. Boxplots show the median, 25^th^ and 75^th^ percentiles, and outliers; black dots indicate individual cells. **(g)** Percentage of miRNA-mapped reads in small-RNA-seq libraries from mouse oocytes and early embryos^28^. Dots represent replicate samples; boxplots show the group mean (vertical line) ± standard deviation (whiskers). **(h)** Log-transformed, mean TMM-normalized expression of miRNA seed families (colored) in mouse oocytes and embryos^28^, showing the top five families, fam-92, and fam-290 (orthologous to human fam-372; top), and the contribution (percentage of total family expression) of individual miRNAs within fam-92 and fam-290 (bottom). Ribbons for fam-92 and miR-290 show the standard deviation across stages (top). **Abbreviations:** (GV) Germinal vesicle oocyte; (MI) Metaphase I oocyte; (MII) Metaphase II oocyte; (1C) one-cell stage embryo; (2C) two-cell stage embryo; (4C) four-cell stage embryo; (8C) eight-cell stage embryo; (EB) embryoid body; (ICM) inner cell mass; (TE) trophectoderm; (fam-92) seed family miR-25-3p/32-5p/92-3p/363-3p/367-3p; (fam-372) seed family miR-302-3p/372-3p/373-3p/520-3p; (fam-290) miR-290 seed family.

We next integrated and analyzed sc-sRNA-seq datasets from Russell et al.^3^, which spans embryos from the 8C through the first differentiated lineages, ICM and TE (**Fig. 1D, Extended Data Fig. 1B**). The proportion of sRNA reads mapping to miRNAs remained relatively stable across the morphology-based and transcriptome-inferred stages and lineages, with ICM cells displaying the highest median miRNA fractions (**Fig. 1D, Extended Data Fig. 1B**). Members of fam-92 and −372 displayed distinct stage-specific patterns of miRNA read proportions and could be categorized into three groups: up-regulated (hsa-miR-367-3p and −302a/b/d-3p, −520c/d-3p), down-regulated (hsa-miR-25-3p, −92a/b-3p), and stably expressed (hsa-miR-32-5p, −363-3p and 372/3-3p, 520a-3p) from 8C to ICM/TE (**Fig. 1E, Extended Data Fig. 1C**). Notably, the highly expressed hsa-miR-92a-3p and hsa-miR-372-3p, displayed modest down-(**Fig. 1F**) and up-regulation (**Extended Data Fig. 1D),** respectively, from 8C onward, suggesting that although both are required at high abundance, their fine-tuned regulation may be important for early lineage differentiation.

To contextualize these findings across species, we re-analyzed bulk sRNA-seq data of mouse embryos^28^. Oocytes and 1-cell stage (1C) embryos segregated distinctly from 2C, 4-cell stage (4C) and 8C embryos in principal component analysis (PCA) (**Extended Data Fig. 1E, Supplementary Tables 5-6**). miRNA-mapped reads increased from the 2C onward, coinciding with mouse major EGA^29^ (**Fig. 1G**) and mirroring the onset of miRNA expression in human embryos. Although the mouse genome contains orthologs of fam-92 and fam-372 (the miR-290 seed family), these were not the most highly expressed families: instead, let-7 family members, known contributors to ICM formation in mice^30^, dominated expression profiles (**Fig. 1H**). Moreover, individual fam-92/-372 members varied markedly in their relative abundance across stages (**Fig. 1H**) contrasting with the more stable expression observed in humans. These species differences support the view that murine models may not fully recapitulate human early-embryo miRNA biology^6^, including the roles of fam-92 and fam-372, and hPSC–based models could provide a more appropriate platform for mechanistic studies.

### miRNA and RNA expression trajectories in naïve hESC-HLC-EXMC conversion

miRNA expression has been profiled in epiblast-resembling naïve and primed hESCs^31,32^ and ICM- and TE-like cells^3,33^, but HLCs and the recently identified EXMCs remain largely unexplored^21–24^. We applied RACL-medium to naïve hESCs to generate a mixed population comprising epiblast-intermediates, HLCs, and EXMCs^21–23^. RACL cells (RACLs) were passaged further in NACL medium to generate expandable EXMCs^21,22^ and we profiled miRNA expression along the naïve hESC-RACL-NACL trajectory. Consistent with reported RNA-level heterogeneity in RACL cultures^22^, immunostaining showed retention of some NANOG-positive (^+^), OCT4^+^, and KLF17^+^ cells, while most cells were GATA6^+^ (early hypoblast marker; **Fig. 2A**). NACL cells (NACLs) were NANOG-negative (^-^), OCT4^-^and KLF17^-^, and uniformly GATA6^+^, suggesting lower heterogeneity. GATA3 was not detected (**Extended Data Fig. 2A**) and FOXA2 (**Fig. 2A)** detection varied between RACL differentiations, suggesting asynchronous or incomplete RACL differentiation at confluence (day 7 (D7) or day 8). At the RNA level, pluripotency markers (*NANOG, KLF17, TFCP2L1, OCT4*) were down-regulated and hypoblast markers (*GATA6, GATA4, SOX17*) up-regulated upon the procession from naïve hESCs to NACLs (**Fig. 2B**). EXM markers (*VIM, HAND1, LUM, POSTN)* were also upregulated, supporting mixed identity in RACLs and an EXMC identity of NACLs. TE markers (*GATA2, GATA3, TEAD3*) were upregulated in RACLs, consistent with a possible TE intermediate in EXMC emergence^22^.

**Figure 2.**
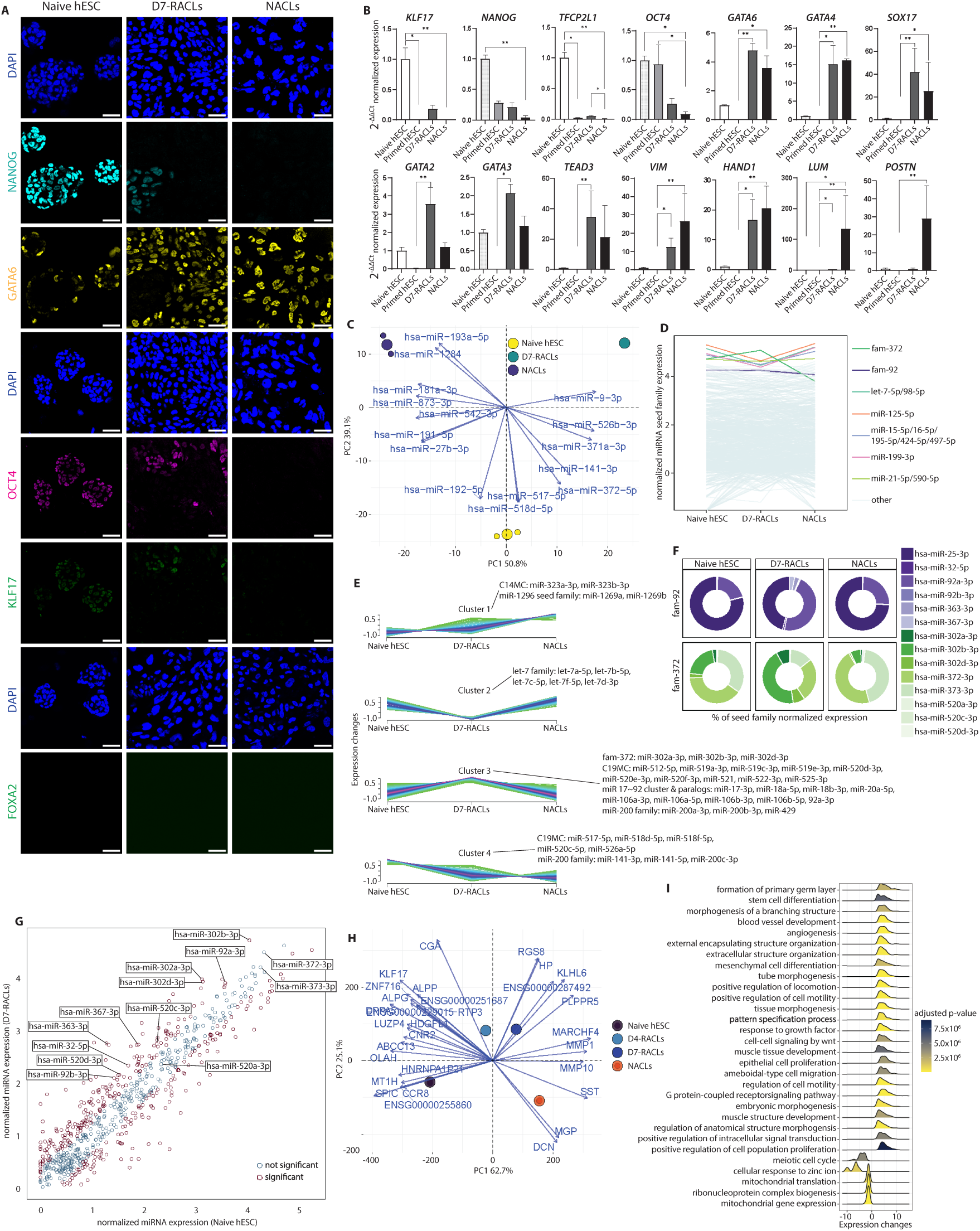
**(a)** Naïve H9 hESCs, D7-RACLs and NACLs immunostained with antibodies recognizing NANOG (cyan), GATA6 (yellow), OCT4 (magenta), KLF17 (green), FOXA2 (green), and DNA counterstained with 4′,6-diamidino-2-phenylindole (DAPI). Scale bars 50 μm. **(b)** Expression levels of pluripotency/epiblast (*KLF17, NANOG, TFCP2L1, OCT4*), hypoblast (*GATA6, GATA4, SOX17*), trophectoderm (*GATA2, GATA3, TEAD3*), and extraembryonic mesoderm (*VIM, HAND1, LUM, POSTN*) marker genes determined by qPCR in primed and naïve H9 hESCs, D7-RACLs and NACLs. The data represent n = 3 replicates shown as mean + standard deviation with the expression levels shown as the difference to the levels in naïve hESCs. Statistical significance was determined using Kruskal-Wallis test followed by Dunn’s test (*p < 0.05, **p < 0.01). **(c)** PCA of log-transformed TMM-normalized expression of all detected miRNAs in naïve H9 hESCs, D7-RACLs, and NACLs (n = 2). Dot color indicates sample type; larger dots represent centroids for the replicates. Vectors indicate top-contributing miRNAs. See Extended Data Figure 2b. **(d)** Log-transformed, mean TMM-normalized expression of miRNA seed families (colored) in naïve H9 hESCs, D7-RACLs, and NACLs, showing the top five families, fam-92 and fam-372. **e)** Fuzzy clustering of miRNA expression in naïve H9 hESCs, D7-RACLs, and NACLs (Supplementary Table 10), with central miRNAs and differential expression (DESeq2: |log₂FC| > 1, FDR < 0.05) status (Supplementary Table 7-9) as follows: Cluster 1: chromosome 14 miRNA cluster (C14MC) members (hsa-miR-323a-3p; trending upregulation in NACLs vs. naïve hESCs or RACLs, hsa-miR-323b-3p: significantly upregulated in NACLs vs. naïve hESCs, trending upregulation in NACLs vs. RACLs) and miR-1296 seed family members (hsa-miR-1269a, −1269b; significantly upregulated in NACLs vs. naïve hESCs or RACLs). Cluster 2: let-7 family members (hsa-let-7a-5p, −7b-5p, −7c-5p, −7f-5p, −7d-3p; significantly downregulated in RACLs vs. naïve hESCs or NACLs). Cluster 3: fam-372 members (hsa-miR-302a-3p, −302b-3p, −302d-3p; significantly upregulated in RACLs vs. naïve hESCs or NACLs), miR-17∼92 cluster and its paralog members (hsa-miR-17-3p, −18a-5p, −18b-3p, −20a-5p, −106a-3p, −106a-5p, −106b-5p, −92a-3p; significantly upregulated in RACLs vs. naïve hESCs or NACLs, hsa-miR-106b-3p: trending upregulation in RACLs vs. naïve H9 hESCs or NACLs), chromosome 19 miRNA cluster (C19MC) members (hsa-miR-512-5p, −519a-3p, −519c-3p, −519e-3p, −520d-3p, −520e-3p, −520f-3p, −521, −522-3p, −525-3p; significantly upregulated in RACLs vs. naïve hESCs or NACLs), and miR-200 family members (hsa-miR-200a-3p, −200b-3p, −429; significantly upregulated in RACLs vs. naïve hESCs or NACLs). Cluster 4: C19MC members (hsa-miR-517-5p, −518f-5p; significantly upregulated naïve hESCs vs. RACLs and trending upregulation in naïve hESCs vs. RACLs, hsa-miR-518d-5p, −520c-5p, −526a-5p; significantly upregulated in naïve hESCs vs. RACLs or NACLs) and miR-200 family members (hsa-miR-141-3p, −141-5p, −200c-3p; significantly upregulated in naïve hESCs vs. RACLs or NACLs). **(f)** Contribution (percentage of total family expression) of individual miRNAs within fam-92 and fam-372 in naïve H9 hESCs, D7-RACLs, and NACLs. **(g)** Scatter plot of log-transformed, mean TMM-normalized miRNA expression in naïve H9 hESCs and D7-RACLs (n = 2). Significantly differentially expressed miRNAs (DESeq2: |log₂FC| > 1, FDR < 0.05) are shown in red; non-significant miRNAs in grey. Members of fam-92 and fam-372 are labeled. **(h)** PCA of log-transformed, TMM-normalized expression of differentially expressed genes (DESeq2: |log₂FC| > 1, FDR < 0.05; Supplementary Tables 11-13) in naïve H9 hESCs, D4-RACLs, D7-RACLs, and NACLs (n = 2). Dot color indicates sample type; larger dots indicate centroids. Vectors indicate top-contributing genes. See Extended Data Figure 3a. **(i)** Enriched biological processes among genes differentially expressed between D7-RACLs and naïve H9 hESCs, filtered for Gene Ontology term hierarchy. Color indicates adjusted p-value; x-axis shows log-fold-change distribution. **Abbreviations:** (D4-RACLs) day 4 RACL cells; (D7-RACLs) day 7 RACL cells; (NACLs) NACL cells; (fam-92) seed family miR-25-3p/32-5p/92-3p/363-3p/367-3p; (fam-372) seed family miR-302-3p/372-3p/373-3p/520-3p.

To define miRNA composition, we performed sRNA-seq on naïve hESCs, RACLs, and NACLs. The replicate samples clustered tightly with RACLs and NACLs flanking naïve hESCs along principal component 1 (PC1) and separating along PC2 (**Extended Data Fig. 2B**). Most detected miRNAs were shared across cell types (**Extended Data Fig. 2C**). Correlation analysis of the miRNA profiles placed RACLs closer to naïve hESCs than NACLs (**Extended Data Fig. 2D**). PCA loadings indicated that hsa-miR-193a-5p, −1284, −181a-3p, and −873-3p contributed strongly to NACLs and hsa-miR-9-3p contributed to RACLs along the first two PCs (**Fig. 2C**). Similarly, hsa-miR-192-5p, −518d-5p, 517-5p, and −372-5p that include established pluripotency and C19MC miRNAs contributed to the naïve hESCs. At the family level, HLCs and EXMCs recapitulated features of human embryos, with high miR-21-5p/590-5p and miR-125-5p abundances (**Fig. 2D**). Contrarily, the let-7 family was also prominent in the hESC-derived samples, consistent with its role in hESC differentiation^34^. Fuzzy clustering^35^ combined with differential expression (DE) analysis (DESeq2)^36^ delineated four miRNA expression trajectories across the naïve hESC-RACL-NACL differentiation: up-regulated, RACL-summit, RACL-peak, and down-regulated miRNAs (clusters 1–4; **Fig. 2E, Supplementary Tables 7-10**). Upregulated miRNAs included C14MC members (hsa-miR-323a-3p, −323b-3p), and the miR-1296 seed family (hsa-miR-1269a, −1269b) among others. RACL-summit miRNAs contained let-7 family members, drivers of pluripotency exit and differentiation^34,37^ that are higher in mouse ICM than TE^30^, suggesting they may restrain HLC and EXMC differentiation. RACL-peak miRNAs included multiple members of fam-372 (hsa-miR-302a-3p, −302b-3p, −302d-3p) and the miR-17∼92 cluster and its paralogs (hsa-miR-17-3p, −18a-5p, −18b-3p, −20a-5p,-106a-3p, −106a-5p, −106b-3p, −106b-5p, −92a-3p; **Fig. 2F-G**), that are ESCC miRNAs^14^. Primate-specific placental C19MC members (hsa-miR-512-5p, −519a-3p, −519c-3p, −519e-3p, −520d-3p, −520e-3p, −520f-3p, −521, −522-3p, −525-3p) and miR-200 family members (hsa-miR-200a-3p, −200b-3p, −429), which regulate EMT^38^, were also among RACL-peak miRNAs. Conversely, down regulated miRNAs included other C19MC (hsa-miR-517-5p, −518d-5p, −518f-5p, −520c-5p, −526a-5p) and miR-200 family members (hsa-miR-141-3p, −141-5p, −200c-3p) indicating intra-cluster and intra-family rewiring between RACLs and NACLs. Within fam-92 and fam-372, hsa-miR-92a-3p and hsa-miR-302b, respectively, reached the highest levels and were significantly upregulated in RACLs (**Fig. 2G, Extended Data Fig. 2E-F**). In naïve hESCs and NACLs, hsa-miR-25-3p (fam-92) as well as hsa-miR-372-3p/-373-3p (fam-372) were most prevalent, with the latter two down-regulated in NACLs. Thus, despite sharing core pluripotency miRNAs, naïve hESCs, HLCs, and EXMCs display distinct miRNA compositions, particularly within C19MC, and hsa-miR-92a-3p and −372-3p are dynamically regulated across RACL and NACL differentiation.

To elucidate cellular programs underlying HLC and EXMC formation, possibly coupled with miRNA regulation, we performed RNA-seq on naïve hESCs, day 4 (D4)-RACLs, D7-RACLs, and NACLs. D4 was included because GATA6^+^ cells emerge around this time and this intermediate state has been under characterized. It may also provide information on the origins of the EXM, which remains to be defined in human embryos but has been studied in other EXMC models and primates^25^. PCA based on DE genes from pairwise comparisons of D4- and D7-RACLs and NACLs against naïve hESC revealed clear clustering of the replicates (**Fig. 2H**). PCA and correlation analysis using all detected transcripts (**Extended Data Fig. 3A-B)** positioned D4- and D7-RACLs close to each other and intermediate between naïve hESCs and NACLs. Top PC loadings of the first two dimensions included naïve markers (*KLF17*^39^*, DPPA5*^39^*, ALPG*^40^) segregating D4-RACLs, and ICM markers (*CCR8*^41^*, SPIC*^42^*, MT1H*^43^) segregating naïve hESCs. Interestingly, the placental hCG-hormone encoding TE marker, *CGA*^44^, contributed to D4-RACL placement along PC1 and PC2. D7-RACLs were associated with endodermal differentiation genes, including pancreatic and hepatic genes (*RGS8*^45^*, HP*^46^). PC1 genes tracking the progression from naïve hESCs to the differentiated states were enriched in ECM and cell migration-related genes (*MARCHF4*^47^*, MMP1, MMP10, MGP*^48^), and the EXMC marker *DCN*^22^, contributed to NACL positioning along the first two PCs.

Gene set enrichment analysis (GSEA) of DE genes (**Supplementary Tables 11-13**) revealed shared biological processes across D4-RACLs, D7-RACLs, and NACLs (**Fig. 2I, Extended Data Fig. 3C**): embryonic differentiation (e.g. formation of primary germ layer, blood vessel development, tissue morphogenesis, response to growth factor), and cell motility and proliferation (positive regulation of locomotion, positive regulation of cell motility, and cell proliferation). D4- and D7-RACL up-regulated processes were also related to differentiation (mesenchymal cell differentiation, pattern specification process) and WNT signaling. D7-RACL and NACL upregulated processes were related to extracellular structure organization, and NACLs displayed muscle tissue related processes. Down-regulated processes in D4- and D7-RACLs included ATP synthesis, mitochondrial transcription and translation, as well as mitotic cell cycle. Analysis of enriched regulatory target gene sets of transcription factors (TFs) and miRNAs, where genes are grouped by their regulatory elements in non-protein coding regions, revealed up-regulation of several GATA TF motifs in D4- and D7-RACLs, as expected (**Extended Data Fig. 3D**). D4 up-regulated motifs included LEF1, a downstream transcription factor of the WNT signaling pathway, as well as MEF2^49^, a key regulator of differentiation of multiple cell types. Up-regulated motifs in D7-RACLs were enriched for Forkhead box (FOX) motifs (FREAC4 01, RTAACA FREAC2 01, FOXH1 TARGET GENES) among others, and a motif for mir-27a/b, which regulates the pluripotency-associated ACTIVIN/NODAL axis (SMAD2/3) of the TGF-ß signaling pathway^50^. Notably, hsa-mir-27a/b-3p were RACL-summit miRNAs (**Supplementary Table 10**), consistent with an inverse relationship between motif activity and miRNA abundance. Together, these analyses indicate that progression toward HLC and EXMC fates is accompanied by extracellular remodeling and coordinated shifts in cell cycle and proliferation, likely orchestrated by pluripotency-associated miRNAs and C19MC members, among others.

### Hsa-Mir-92a-3p is required for timely differentiation of HLCs and EXMCs

Both hsa-miR-92a-3p and −372-3p are established regulators of stem cell identity, and loss of fam-92 family members has been associated with defects in lung, heart, and liver development in mice^51^. Consistent with this, GSEA of naïve hESC-to-EXMC differentiation revealed enrichment of up-regulated terms associated with these tissues, suggesting that hsa-miR-92a-3p may play a functional role in HLC and EXMC differentiations. To test this, we inhibited hsa-miR-92a-3p (92i) using a validated LNA-inhibitor ^52,53^, alongside a seed-mismatch control (92MM) and a non-targeting scramble (NegA) during RACL differentiation. LNAs were delivered via gymnosis from day 1 to D4 or D7 (**Fig. 3A**). Initial titrations (0.1–3 µM) revealed increasing toxicity as the concentration approached 3 µM (data not shown). Therefore, we used concentrations up to 0.75 µM to achieve inhibition without compromising viability. Because LNA treatment does not necessarily reduce mature miRNA levels, we validated inhibition indirectly by quantifying the known hsa-miR-92a-3p target *CDKN1C*^54,55^. *CDKN1C* expression increased on D7 in 92i cells in a dose-dependent manner (**Fig. 3B, Extended Data Fig. 4A**). Internalization of the inhibitor was confirmed using a 5’ fluorescein (FAM)-labeled LNA and confocal microscopy (**Extended Data Fig. 4B**). RNA-seq of D4- and D7-RACLs, together with naïve and primed hESCs and NACLs, revealed that 92i samples clustered within the RACL group and displayed good replicate overlap (**Fig. 3C; Extended Data Fig. 4C**). Notably, 92i D7-RACLs (0.5 and 0.75 µM) shifted toward D4-RACLs along PC1, suggesting a differentiation delay. Integration with human embryo scRNA-seq data^23,56^ confirmed lagging progression along the epiblast-to-hypoblast trajectory normally followed by the naïve hESC-RACL-NACL sequence (**Fig. 3D; Extended Data Fig. 4D**).

**Figure 3.**
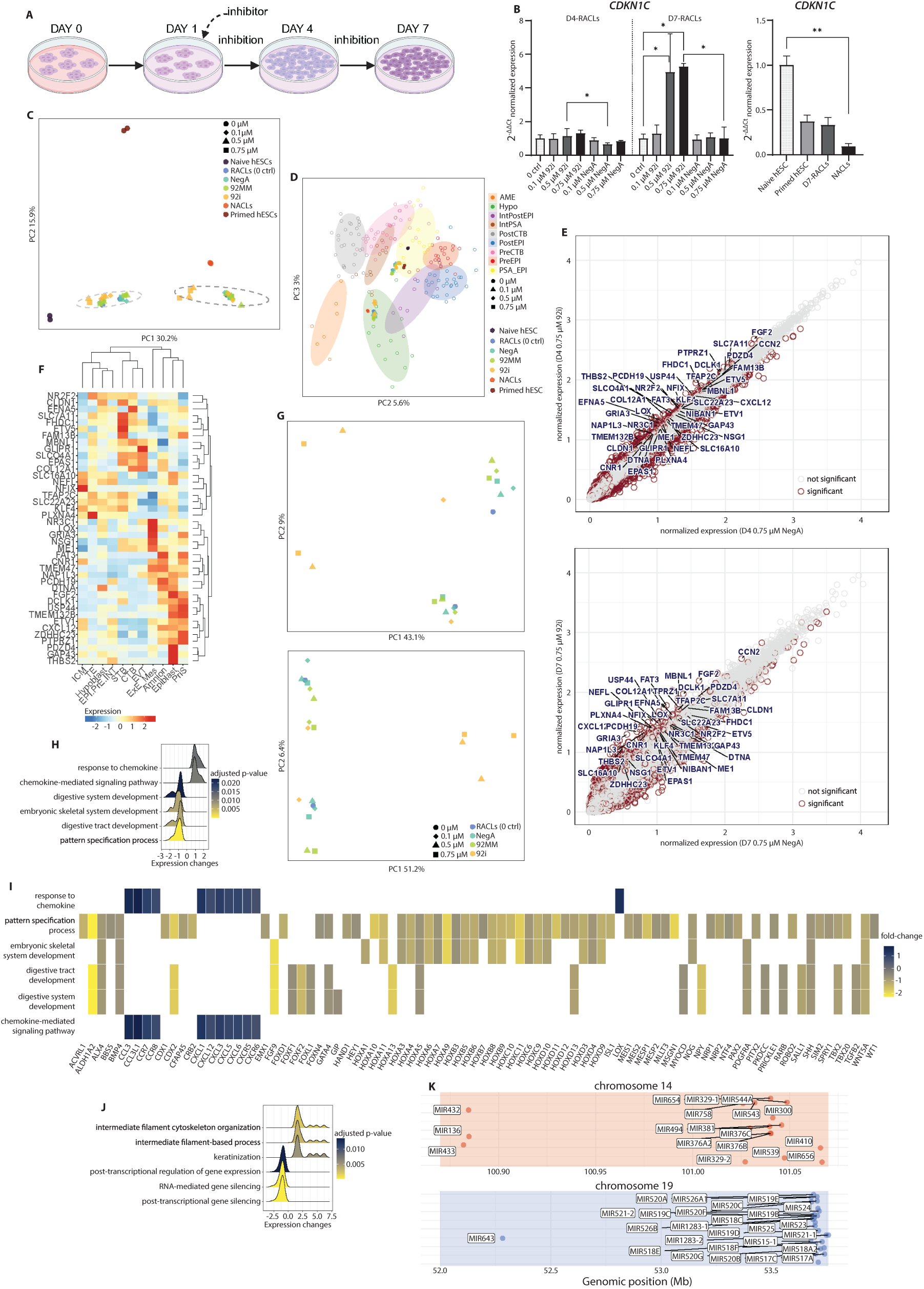
**(a)** Schematic of LNA inhibitor or control treatment and sample-collection timepoints during RACL differentiation of naïve H9 hESCs. **(b)** *CDKN1C* expression measured by qPCR in D4-and D7-RACLs that were untreated (0 ctrl) or treated with 92i or NegA at the indicated doses (left), and in primed and naïve H9 hESCs, D7-RACLs, and NACLs (right). The data represent n = 2 (left panel) and n = 3 (right panel) replicates shown as mean + standard deviation. Statistical significance was determined using Kruskal-Wallis test followed by Dunn’s test (*p < 0.05, **p < 0.01). **(c)** PCA of log-transformed TMM-normalized gene expression in primed and naïve H9 hESCs, NACLs, and D4- and D7-RACLs that were untreated (0 ctrl) or treated with 92i, 92MM, or NegA (n=2). Shapes indicate treatment dose; colors indicate cell type or treatment group for the RACL samples. See Extended Data Figure 4c. **(d)** PCA (PC2 and PC3) of RNA-seq data from this study (samples as in panel c) together with human embryo single cell RNA-seq data^56^, annotated according to Okubo et al.^23^ Colored ovals indicate embryonic cell-type clusters with individual datapoints shown as colored rings: (PreEPI) pre-implantation epiblast; (PostEPI) post-implantation epiblast; (PSA_EPI) primitive-streak anlage epiblast; (IntPSA and IntPostEPI) intermediate state cells of primitive-streak anlage epiblast and post-implantation epiblast; (AME) amnion; (Hypo) hypoblast; (preCTB) pre-implantation trophectoderm; (postCTB) post-implantation cytotrophoblast. **(e)** Scatter plots of log-transformed, mean TMM-normalized gene expression in D4-RACLs (n = 2; top) and D7-RACLs (n = 2; bottom) treated with 0.75 μM 92i or NegA. Highly expressed (mean expression in the considered samples > 600) and significantly up-regulated predicted hsa-miR-92a-3p targets (TarBase)^106^ are labeled. Significantly differentially expressed genes (see methods for sample groups; DESeq2: |log₂FC| > 0.5, FDR < 0.05) are shown in red; non-significant genes in grey. **(f)** Clustered heatmap showing the expression of the labeled hsa-miR-92a-3p targets from panel (e) in human embryo single cell RNA-seq data^57^ using the Human Embryonic Reference Tool (v1.1.1.17). Abbreviations: (ICM) inner cell mass; (TE) trophectoderm; (EPI.PrE.INT) epiblast–hypoblast intermediates; (STB) syncytiotrophoblast; (CTB) cytotrophoblast; (EVT) extravillous trophoblast; (ExE_Mes) extraembryonic mesoderm; (PriS) primitive streak. **(g)** PCA of differentially expressed genes between 92i and control samples for D4-RACLs (top) and D7-RACLs (bottom; see methods for sample groups; DESeq2: |log₂FC| > 0.5 and FDR < 0.05; Supplementary table 14). Colors and shapes indicate RACL sample type as in panel (c). **(h)** Enriched biological processes among differentially expressed genes between 92i and control D4-RACLs filtered for Gene Ontology term hierarchy. Color indicates adjusted p-value; x-axis shows log-fold-change distribution. **(i)** Differentially expressed genes between 92i and control D4-RACLs included in the enriched biological processes shown in panel (h). Color key indicates fold-change values. **(j)** Enriched biological processes among differentially expressed genes between 92i and control D7-RACLs, filtered for Gene Ontology term hierarchy. Colors indicate adjusted p-values; x-axis shows log-fold-change distribution. **(k)** Genomic locations of chromosome 14 miRNA cluster (C14MC) and chromosome 19 miRNA cluster (C19MC) members included among the RNA-mediated gene-silencing–related biological processes enriched in D7-RACLs (see panel (j)). **Abbreviations:** (D4-RACLs) day 4 RACL cells; (D7-RACLs) day 7 RACL cells; (NACLs) NACL cells; (92i) hsa-miR-92a-3p inhibitor; (92MM) seed-mismatch control; (NegA) non-targeting control.

To identify dysregulated hsa-miR-92a-3p targets, we performed DE analysis for D4 and D7 separately, comparing 92i (0.5, 0.75 µM) against grouped controls (medium, NegA, 0.1 µM 92i; **Supplementary Table 14**), as determined by PCA. We focused on genes up-regulated at both D4 and D7, consistent with miRNA inhibition reducing target mRNA decay (**Extended Data Fig. 4E**), and required validated 3′UTR binding by hsa-miR-92a-3p based on TarBase^54^. Potential targets showed no expression differences between 0.1 uM 92i and NegA samples on D4 and D7, as expected (**Extended Data Fig. 4F)**. Highly expressed up-regulated targets included naïve and primed markers (*FGF2, KLF4, USP44*) and TE-associated genes (*TFAP2C, CXCL12, NR2F2*) (**Fig. 3E**). Because miRNAs can also act outside 3′UTRs^1^, we examined additional up-regulated genes that may represent alternative targets or non-targets related to phenotypic differences. These included the pluripotency and TE markers, *SOX2* and *CGA*, respectively (**Extended Data Fig. 4G**). Mapping hsa-miR-92a-3p targets onto human embryo scRNA-seq data^57^ revealed three major expression clusters: a small cluster enriched in early embryonic states (ICM, TE, hypoblast, epiblast, epiblast–hypoblast intermediates), and two larger clusters enriched in advanced TE derivatives (cytotrophoblast, syncytiotrophoblast (STB), and extravillous trophoblast (EVT)) and later embryonic, especially epiblast-derived states (extraembryonic mesoderm, amnion, epiblast, and primitive streak)^58^ (**Fig. 3F**). This indicates that many derepressed targets in 92i RACLs are associated with epiblast or TE fates, supporting that hsa-miR-92a-3p inhibition impedes exit from the epiblast-like naïve state. PCA of all DE genes from D4 and D7 comparisons showed clear separation between 92i and control samples along PC1, with a minor batch effect along PC2 (**Fig. 3G**). GSEA of D4 DE genes revealed up-regulation of chemokine signaling and down-regulation of embryonic differentiation pathways in 92i RACLs (**Fig. 3H**). CC- and CXC-class chemokines were strongly up-regulated upon 92i, consistent with a TE-like shift within the extra-embryonic RACL differentiation context (**Fig. 3I**)^59^. Conversely, multiple FOX and homeobox (HOX) family developmental regulators^60^, as well as hypoblast-related genes (*GATA4, PDGFRA*), were down-regulated. Up-regulated genes in 92i D7-RACLs were related to cytoskeletal organization (**Fig. 3J**), including multiple keratin genes (**Extended Data Fig. 4H**). Terms related to RNA-mediated gene silencing were down-regulated, and many affected genes were miRNA or miRNA host genes. Among these were C14MC and C19MC miRNAs, associated with ICM or post-implantation trophoblast cells^61^ and TE, respectively^3^ and dynamically regulated during the RACL/NACL differentiation (**Fig. 3K**). Target motif-based GSEA identified several miRNA motifs upregulated in 92i D7-RACLs, while downregulated motifs included LEF1, and HOXA13, the latter of which is enriched in highly accessible chromatin of EXMCs^22^ (**Extended Data Fig. 4I**). Together, these findings strongly indicate that hsa-miR-92a-3p inhibition disrupts RACL differentiation, producing a dysregulation of multiple target genes and developmental regulators, and delaying the transition from an epiblast-like state toward hypoblast and EXM fates.

### FGF2 signaling is dysregulated upon hsa-miR-92a-3p inhibition

*FGF2* emerged among the most highly expressed, validated targets of hsa-miR-92a-3p and FGF signaling is known to support human hypoblast differentiation^62,63^, while FGF2 maintains primed hESC identity^64^. We therefore investigated FGF signaling dynamics during RACL differentiation and in response to 92i (**Fig. 4A**). In on our RNA-seq data, *FGF2* expression was lowest in naïve hESCs and NACLs, increased at D4 and decreased again at D7 in RACLs. In contrast, 92i elevated *FGF2* expression at both D4 and D7, with D7 levels resembling those of primed hESCs. *FGF4*, a naïve hESC marker^39^, was highly expressed at D4 and declined at D7, with slightly elevated levels in 92i RACLs. *FGFR1* increased from naïve hESCs through D4- and D7-RACLs and decreased in NACLs, with minimal differences between 92i and controls. *FGFR2* expression was up-regulated only at D7, showed a modest reduction in 92i RACLs, and reached its highest levels in NACLs. Trajectory inference using Slingshot^65^ and mean pseudotime analysis of D7-RACL and NACL scRNA-seq data^22^ further supported a structured sequence of FGF/FGFR dynamics across the two-forked differentiation trajectory from a epiblast-intermediate state towards HCLs and EXMCs (**Fig. 4B**). *FGF2, FGF4, FGFR3*, and *FGFR4* peaked in early pseudotime within epiblast-intermediates (**Fig. 4C**). *FGFR1* seemed to precede *FGFR2* during later pseudotime in RACL-derived HLCs and EXMCs and in NACL EXMCs. This pattern mirrors mouse hypoblast differentiation, in which NANOG⁺ ICM cells produce FGF ligands and GATA6⁺ hypoblast precursors transition from FGFR1 to FGFR2 expression during fate commitment ^66,67^.

**Figure 4.**
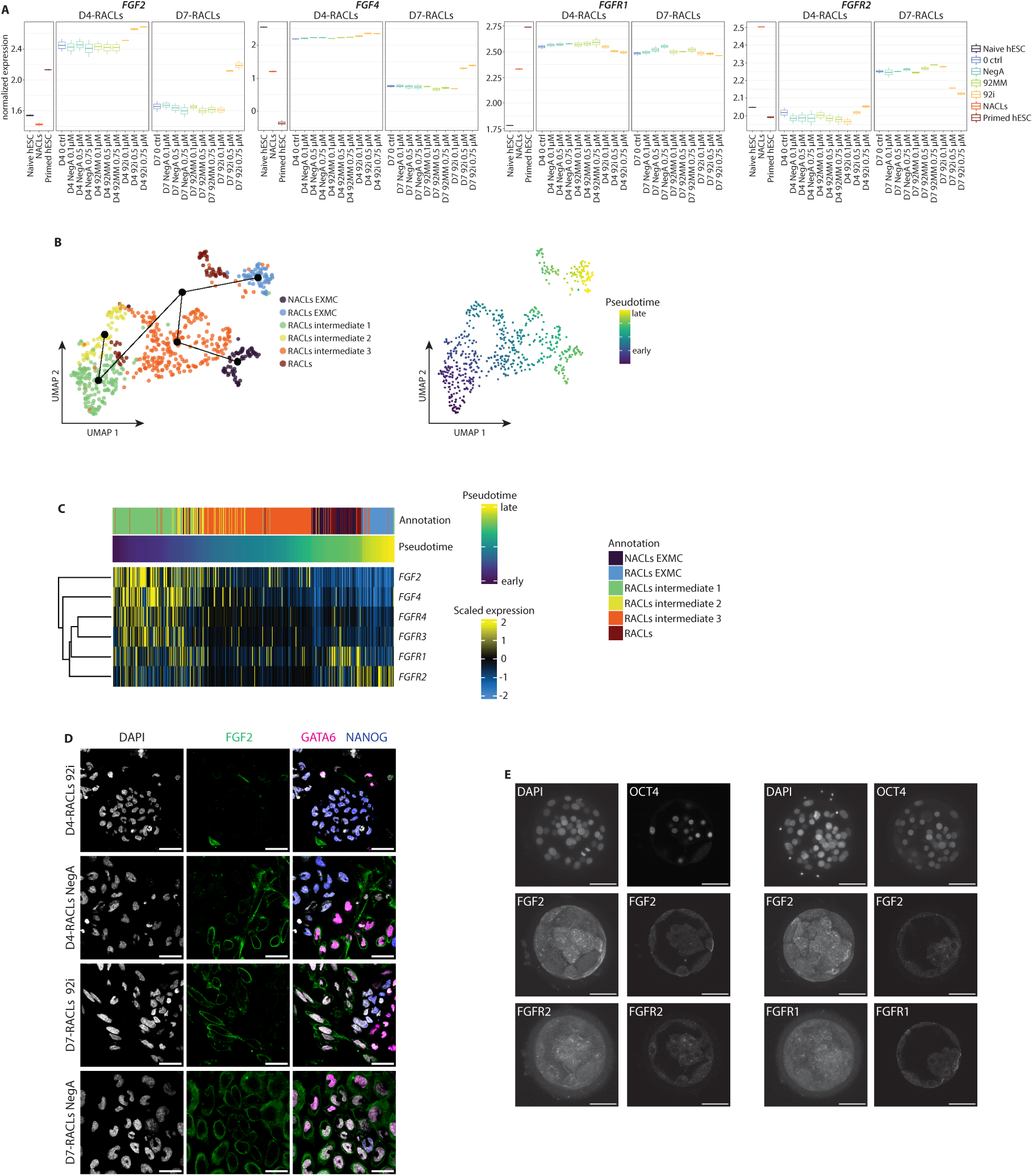
**(a)** Log-transformed, mean TMM-normalized expression levels of *FGFs* and *FGFRs* measured by RNA-seq in D4-and D7-RACLs that were untreated (0 ctrl) or treated with 92i, 92MM, or NegA at the indicated doses, as well as in primed and naïve H9 hESCs and NACLs. Cell type or treatment group for the RACL samples (n=2) is indicated in colors. The boxplot indicates the median, the 25^th^ and 75^th^ percentiles. **(b)** UMAP of cell type annotations from Pham et al.^22^ overlayed with the Slingshot-inferred^65^ lineage trajectory (left) and mean pseudotime (right) of D7-RACL and NACL single cell RNA-seq data^22^. **(c)** *FGF* and *FGFR* expression in D7-RACLs and NACLs^22^ ordered according to mean pseudotime with scaled normalized expression shown. Colors indicate cell type annotations by Pham et al.^22^ Rows were clustered using Euclidean distance and average linkage. **(d)** D4- and D7-RACLs treated with 92i or NegA (0.75 μM) immunostained with antibodies recognizing FGF2 (green), GATA6 (magenta), NANOG (blue), and DNA counterstained with 4′,6-diamidino-2-phenylindole (DAPI). Scale bars 40 μm. **(e)** Day 6 post fertilization human blastocysts immunostained with antibodies recognizing OCT4, FGF2, FGFR1, and FGFR2. DNA counterstained with 4′,6-diamidino-2-phenylindole (DAPI). Scale bars 50 μm. Representative images shown (n=2). **Abbreviations:** (D4-RACLs) day 4 RACL cells; (D7-RACLs) day 7 RACL cells; (NACLs) NACL cells; (92i) hsa-miR-92a-3p inhibitor; (92MM) seed-mismatch control; (NegA) non-targeting control.

We next assessed FGF signaling at the protein level. Immunostaining for FGF2, FGFR1, and FGFR2 in naïve hESCs, and D4- and D7-RACLs (92i and NegA controls), together with epiblast (NANOG, SUSD2) and hypoblast (GATA6) markers, revealed multiple regulatory defects. As expected, naïve hESCs showed low FGF2 levels (**Extended Data Fig. 5A**). At D4, FGF2 signal localized at the cell membrane of GATA6⁺ cells in controls, whereas FGF2⁺/GATA6⁺ cells were infrequent in 92i RACLs (**Fig. 4D**). By D7, controls showed robust FGF2 co-staining with low- to high-intensity GATA6⁺ cells. A comparable but markedly reduced signal was observed in 92i RACLs, which retained more NANOG⁺ cells than controls, consistent with impaired exit from an epiblast-like state. These defects may reflect insufficient FGF2 production or impaired ligand recruitment via FGFRs. FGFR1 signal was low in naïve hESCs and upregulated at D4 in both conditions, without obvious co-staining preference for NANOG or GATA6 (**Extended Data Fig. 5B**). By D7, controls were mainly GATA6⁺/FGFR1⁺, whereas this population was reduced in 92i RACLs. FGFR2 staining was observed at the cell membrane in naïve hESCs, and cytoplasmic and focal staining was observed at D4 and D7, with no major differences between conditions (**Extended Data Fig. 5C**). Furthermore, we noted a trending decline of GATA6⁺/NANOG⁻ cells in 92i D7-RACLs (**Extended Data Fig. 5D**). To extend these findings to human embryos, we examined FGF protein localization in blastocysts (Day 6 post fertilization). FGF2 was detectable and appeared stronger in TE than ICM (**Fig. 4E**). FGFR1 and FGFR2 were also present without a conclusive lineage bias. These observations underscore the relevance of FGF2-mediated signaling in early human embryos and suggest that even modest perturbations in this pathway, whether through direct targeting by hsa-miR-92a-3p or secondary effects, may disrupt hypoblast and EXM differentiation.

### Hsa-miR-92a-3p inhibition reduces HLC and EXMC differentiation and blastoid formation

Because the FGF stainings revealed heterogeneity in 92i RACLs, we next examined cell identity more comprehensively. We used lineage-specific marker genes relevant to RACL/NACL differentiation in our RNA-seq dataset (**Fig. 5A**), together with pseudotime ordered D7-RACL and NACL scRNA-seq data (**Fig. 5B**). Across pseudotime, early cells expressed naïve markers (*NANOG*, *KLF17*, *DPPA3*, *TFCP2L1*) and primed markers (*DNMT3B*, *POU5F1*) at high levels (**Fig. 5B**). Notably, several TE markers (*CGA*, *KRT18*, *KRT7*) peaked in mid-pseudotime, particularly in epiblast-intermediate populations (intermediate-3 cells)^22^. Consistently, D4-RACLs exhibited elevated naïve, epiblast, and STB marker expression (**Fig. 5A**). Late pseudotime cells expressed hypoblast (*GATA6, GATA4, BMP2, FOXA2*), amnion (*ISL1, BMP4*), formative (*OTX2*), TE (*GATA2, GATA3, TEAD3*), and EXM (*VIM, NID2, LUM*) markers at high levels (**Fig. 5B**), paralleling the rise of hypoblast and EXM markers in D7-RACLs (**Fig. 5A**). Across D4 to D7, expression of EVT, amnion, primed hESC, trophoblast stem cell (TSC), and core pluripotency markers shifted (**Fig. 5A**). However, 92i D4-RACLs retained slightly higher naïve, primed, epiblast, and STB marker expression than controls. By D7, 92i led to lower EXMC and EVT marker expression and differences in TSC, STB, and naïve marker expression compared to controls. These results were strongly supported by the protein levels: 92i D7-RACLs contained fewer GATA6⁺/GATA4⁺/VIM⁺ cells relative to NegA controls, which were predominantly triple-positive (**Fig. 5C**). In naïve hESCs, only GATA6⁺ cells were present, as expected (**Extended Data Fig. 6A**). Quantification of GATA6^+^ and GATA4^+^ cells in 92i and control D4- and D7-RACLs further confirmed this result (**Extended Data Fig. 6A**). Together, reduced hypoblast and EXM markers and persistence of naïve and TE signatures in both RNA and protein analyses indicate that 92i impedes exit from epiblast-resembling states and biases cells toward earlier RACL differentiation or TE-like identities.

**Figure 5.**
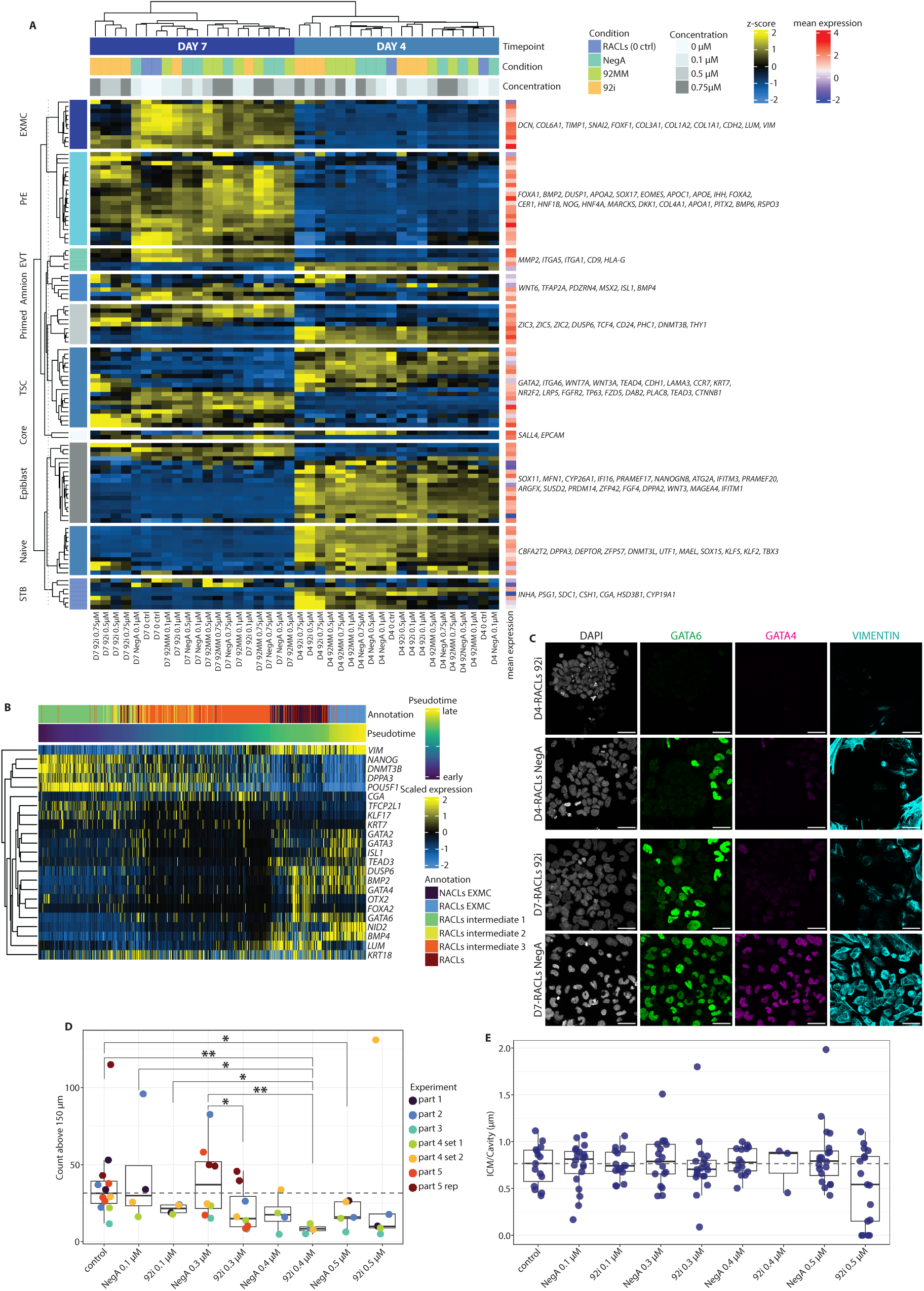
**(a)** Hierarchically clustered expression heatmap (Euclidean distance, average linkage) of lineage-specific marker genes^21^ in primed and naïve H9 hESCs, NACLs, and D4- and D7-RACLs that were untreated (0 ctrl) or treated with 92i, NegA, or 92MM (n=2). “Condition” and “concentration” color keys indicate treatment type and dose. Data are presented as row z-scores of mean TMM-normalized expression. “Mean expression” indicates the mean expression across samples. **(b)** Scaled normalized expression of lineage markers in D7-RACL and NACL single cell RNA-seq data^22^, ordered by mean pseudotime. Marker sets: naïve (*NANOG, KLF17, DPPA3, TFCP2L1*); primed (*DNMT3B, POU5F1*); formative (*OTX2*); trophectoderm (*GATA2, GATA3, TEAD3, CGA, KRT18, KRT7*); hypoblast (*GATA6, GATA4, BMP2, FOXA2*); amnion (*ISL1, BMP4*); extraembryonic mesoderm (*VIM, NID2, LUM*). Colors indicate Pham et al.^22^ cell-type annotations. Rows were clustered using Euclidean distance and average linkage. **(c)** D4- and D7-RACLs treated with 92i or NegA (0.75 μM) immunostained with antibodies recognizing GATA6 (green), GATA4 (magenta), vimentin (cyan), and DNA counterstained with 4′,6-diamidino-2-phenylindole (DAPI). Scale bars 40 μm. **(d)** Number of blastoids formed from naïve H9 hESCs treated during differentiation with NegA or 92i at the indicated concentrations. Boxplots show the median, and 25^th^ and 75^th^ percentiles; dots represent individual blastoids, colored by experiment. The dotted line shows the median of the untreated control. Statistical significance was determined using pairwise Wilcoxon rank-sum tests, with significant p-values indicated in the plot (*p < 0.05, **p < 0.01). **(e)** Inner cell mass (ICM)–to–cavity ratio of blastoids formed from naïve H9 hESCs treated with NegA or 92i during differentiation at the indicated concentrations. The boxplot indicates the median, the 25^th^ and 75^th^ percentiles; dots represent individual blastoids. The dotted line shows the median of the untreated control. No statistically significant differences were observed between groups using pairwise Wilcoxon rank-sum tests. **Abbreviations:** (D4-RACLs) day 4 RACL cells; (D7-RACLs) day 7 RACL cells; (NACLs) NACL cells; (92i) hsa-miR-92a-3p inhibitor; (92MM) seed-mismatch control; (NegA) non-targeting control.

To test whether hsa-miR-92a-3p also regulates extra-embryonic differentiation in a 3D context, we used human blastoids. qPCR confirmed the presence of hsa-miR-92a-3p in blastoids at comparable levels to naïve hESCs (**Extended Data Fig. 6B**). We applied increasing concentrations of 92i and quantified blastoid formation (see Methods). High concentrations (0.4–0.5 µM) impaired blastoid formation even in NegA controls, whereas lower doses yielded comparable blastoid numbers across medium only, NegA, and 92i samples (**Fig. 5D**). At 0.1 µM and 0.3 µM, NegA samples did not differ from medium controls (p = 0.855 and p = 0.908, respectively), while 92i samples showed a decreasing trend in formation (0.1 µM: p = 0.060; 0.3 µM: p = 0.076), suggesting modestly compromised blastoid development. To evaluate lineage composition, we quantified the ICM-to-cavity ratio in a random subset of successfully cavitated blastoids but observed no significant differences between the conditions (**Fig. 5E**). Blastoids from all groups formed embryonic and extra-embryonic compartments, contained a cavity, and reached appropriate diameters resembling blastocysts^68^ (**Extended Data Fig. 6C**), indicating that although 92i reduces blastoid formation efficiency, it does not completely block blastoid development at the used concentrations. These findings mirror the partial differentiation delay observed in RACLs.

### Hsa-miR-92a-3p inhibition perturbs cell-cycle dynamics

Since several miR-17∼92 cluster members are ESCC miRNAs, we next investigated cell-cycle dynamics in naïve hESCs, RACLs, and NACLS. The ESC cell cycle is characterized by a short G1 phase and a high proportion of cells in S phase^69,70^. Because ERK signaling, downstream of FGF, promotes the G1/S transition^71–73^, we asked whether the delayed differentiation in 92i RACLs involves altered cell-cycle regulation. Seurat cell-cycle scoring^74^ of D7-RACL and NACL scRNA-seq data revealed that most NACL EXMCs were in G1, whereas D7-RACL HLCs and EXMCs were in S or G2/M (**Extended Data Fig. 7A**). This indicates that hypoblast and EXM identities may be associated with distinct proliferative states. Consistent with this, Kyoto Encyclopedia of Genes and Genomes (KEGG) cell-cycle genes displayed clear pseudotime-dependent differences: the RACL epiblast-intermediate population 2 and 3 showed a distinct expression profile relative to earlier intermediates and terminal HLC and EXMC states (**Fig. 6A**).

**Figure 6.**
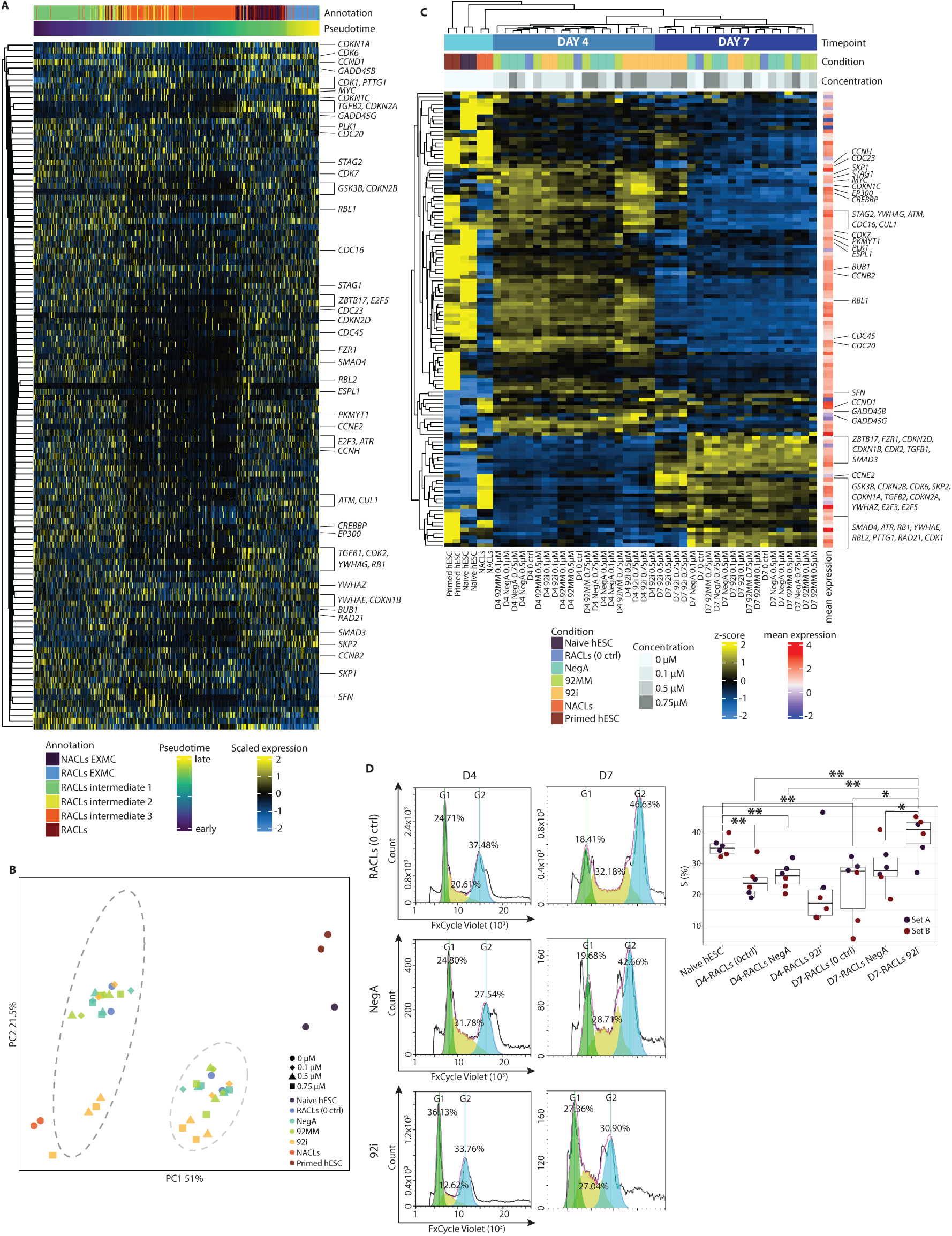
**(a)** Scaled normalized expression of KEGG cell-cycle genes in D7-RACL and NACL single cell RNA-seq data^22^, ordered by mean pseudotime. Colors indicate Pham et al.^22^ cell-type annotations. Rows were clustered using Euclidean distance and average linkage. **(b)** PCA of log-transformed, TMM normalized expression of Seurat’s cell-cycle genes in primed and naïve H9 hESCs, and NACLs, and D4- and D7-RACLs that were untreated (0 ctrl) or treated with 92i, 92MM, or NegA (n=2). Shapes indicate treatment dose; colors indicate cell type or treatment group for the RACL samples. Dotted ovals indicate D4-(light grey) and D7-RACLs (dark grey). **(c)** Hierarchically clustered heatmap (Euclidean distance, average linkage) of KEGG cell-cycle gene expression in primed and naïve H9 hESCs, NACLs, and D4- and D7-RACLs that were untreated (0 ctrl) or treated with 92i, NegA, or 92MM (n=2). “Condition” and “concentration” color keys indicate treatment type and dose. Data are presented as row z-scores of mean TMM-normalized expression. “Mean expression” indicates the mean expression across samples. **(d)** Cell-cycle profiles in D4- and D7-RACLs that were untreated (0 ctrl) or treated with NegA or 92i (0.75 μM; left). DNA content was measured using FxCycle Violet and flow cytometry. Representative data are shown from two differentiation experiments (see Figure S7d for biological replicate) with three technical replicates per sample type (day and treatment). Percentage of S-phase cells in naïve H9 hESCs (see Figure S7c) and RACL subgroups (right). The boxplot indicates the median, the 25^th^ and 75^th^ percentiles; dots represent replicates, colored by differentiation batch. Statistical significance was determined using pairwise Wilcoxon rank-sum tests, with significant p-values indicated in the plot (*p < 0.05, **p < 0.01). **Abbreviations:** (D4-RACLs) day 4 RACL cells; (D7-RACLs) day 7 RACL cells; (NACLs) NACL cells; (92i) hsa-miR-92a-3p inhibitor; (92MM) seed-mismatch control; (NegA) non-targeting control.

PCA of our RNA-seq samples using Seurat’s cell-cycle genes further supported these trends (**Fig. 6B**). The centroids of 92MM, NegA, and medium controls overlapped, but 92i RACLs, shifted away from these groups along PC2. Notably, 92i RACLs diverged from other conditions already at D4, and this separation increased at D7. While control D7-RACLs moved back toward naïve hESCs along PC2, 92i D7-RACLs clustered near the D4 samples and a minor batch effect present at D4 was no longer detectable by D7 (**Extended Data Fig. 7B**). Expression of KEGG cell-cycle genes revealed that naïve and primed hESCs and NACLs were distinct from RACLs, while D4- and D7-RACLs differed substantially from one another (**Fig. 6C**). Within each timepoint, 92i samples formed separate subclusters, yet they still most closely resembled other samples from the same timepoint, indicating modest perturbation of cell-cycle progression rather than a complete rewiring. To validate transcriptome-based predictions, we performed cell-cycle analysis using flow cytometry. Naïve hESCs, and D4- and D7-RACL controls showed no change in G1, an increase in the G2 fraction at D7, and a decline in S beginning at D4 and maintained at D7 (**Fig. 6D, Extended Data Fig. 7C**), confirming that cell-cycle profiles shift across the naïve hESC to HLC/EXMC transition. At D4, 92i and NegA RACLs were comparable. However, fewer 92i D7-RACLS were in G2 and more in S compared to NegA, indicating a delay in G2/M entry and progression. Together, these findings suggest that cell-cycle control contributes to the hsa-miR-92a-3p–dependent differentiation defect. Accurate regulation of hsa-miR-92a-3p is therefore suggested to be critical in coordinating cell-cycle phase transitions with timely hypoblast and EXM specification in early human development.

## DISCUSSION

Our comparative embryo analyses indicate that fam-92 and fam-372 miRNAs are among the most abundant families in early human embryos and are markedly enriched relative to mouse. This highlights their importance not only in the well-established ESC context^14^ but also in early human embryogenesis. In naïve hESCs and in RACL and NACL cultures, which, as shown here and by others^22^, contain epiblast-intermediates, HLCs, EXMCs, and a purer EXMC population, respectively, members of fam-372 and fam-92 were again prominently expressed. Multiple miRNA seed families and clusters were dynamically regulated along the naïve hESC-RACL-NACL trajectory. Notably, several miR-200 family members were highly expressed in naïve hESCs and RACLs, consistent with their involvement in EMT associated transcriptional networks^75^ and with the established contribution of EMT programs to EXMC formation in vitro, suggested by our findings and those of Pham et al.^22^ The miR-200 family is regulated by TGF β/Activin A/Nodal–Smad signaling^13,76^, and Activin A is a component of both RACL and NACL media^21^. Forced expression of miR-200 family members is sufficient to block TGF β-induced EMT, whereas cells that undergo EMT in response to TGF-β downregulate miR-200^75^. Moreover miR-200c, is implicated in hESC renewal and first lineage differentiations, mediated by GATA4 targeting^76^. Together, these observations suggest that downregulation of the miR-200 seed family may be necessary for EXMC formation in both RACL and NACL conditions. Conversely, NACL-derived EXMCs expressed multiple miR-1269 seed family members, which participate in a positive feedback loop with TGF-β signaling: TGF-β induces miR-1269 via SOX4, and miR-1269a further amplifies TGF-β signaling by targeting SMAD7 and HOXD10^77^. This interplay underscores the integration of miRNA regulation with TGF-β pathway dynamics during EXMC generation. Interestingly, RACLs exhibited low expression of let-7 family miRNAs, which are typically associated with promoting pluripotency exit in hESCs^37^. RACLs downregulate canonical pluripotency factors, and the reduced let-7 expression may reflect the extra-embryonic nature of these cultures or the presence of TE-like intermediates. Supporting this interpretation, forced expression of let-7a in early mouse embryos reduces *Cdx2* expression in the TE and causes developmental arrest at the morula stage, when the first lineage decisions have initiated^30^. Moreover, let-7 targets *Tead4*^78^, a key mediator of Hippo-dependent activation of TE genes, including *Cdx2*, in outer morula cells^79,80^. Taken together, these data indicate that miRNAs are intricately coupled with lineage-defining signaling pathways, particularly TGF-β and Hippo-associated circuits, to direct the generation of HLCs and EXMCs during RACL and NACL differentiations.

The C19MC locus showed striking and dynamic regulation across the naïve hESC-RACL-NACL differentiation. This primate-specific cluster, which encodes more than 50 mature miRNAs, is thought to have evolved relatively recently from the smaller ancestral hsa-miR-371∼373 cluster, which has orthologs throughout mammals^81,82^. In naïve hESCs and RACLs, *AAGUGCU*-seed ESCC miRNAs are abundant, but we observed a marked shift in the expression of closely related C19MC miRNAs that are also a hallmark of naïve hESCs, and disruption of this cluster has been shown to hinder the primed-to-naïve conversion^83^. Naïve hESCs preferentially express the 5p arms of several C19MC members (hsa-miR-517-5p, −518d/f-5p, −520c-5p, −526a-5p), which share a conserved seed distinct from the conserved 3p seed. By contrast, several 3p-arm C19MC miRNAs, including the fam-372 member miR-520-3p family (hsa-miR-520d/e/f-3p) and members of the miR-519-3p subfamily (hsa-miR-519a/c-3p) that have a shortened version of *AAGUGCU*-seed^84^, peaked in RACLs. Additional C19MC species with subtly different seed compositions (hsa-miR-512-5p, −519e-3p, −521, −522-3p, −525-3p) also showed elevated abundance, emphasizing that ESCC miRNAs and their evolutionary relatives within C19MC are prominent in naïve and mixed epiblast-intermediate or HLC states but decrease in NACL-derived EXMCs. The downregulation of these early-embryonic C19MC miRNAs may be an essential prerequisite for permitting differentiation into peri- and post-implantation cell types, whose miRNA landscapes remain largely unmapped.

Interestingly, the miR-17∼92 cluster, which appears to have evolved independently of both C19MC and the hsa-miR-371∼373 cluster, also contains ESCC miRNAs with seed sequences similar to fam-372 ESCC miRNAs and several C19MC 3p-arm miRNAs^14^. Members of fam-92, which carry a different seed, are likewise encoded by the miR-17∼92 cluster and its paralogs, further expanding the repertoire of miRNA–mRNA regulatory interactions shared between early human embryos and their ESC-based models. Our integrated analyses position hsa-miR-92a-3p as a rate-limiting regulator that facilitates the exit from epiblast-like states and enables the acquisition of hypoblast and EXM identities during RACL differentiation. Two lines of evidence support this conclusion. First, functional inhibition of hsa-miR-92a-3p with a validated LNA inhibitor^52,53^, led to delayed differentiation characterized by persistent naïve, epiblast, and TE-like transcriptional features and reduced expression of hypoblast and EXM markers. While off-target effects of the LNA cannot be fully excluded, these findings align with prior evidence that miR-92 function is required in vivo. In mice, miR-92 knockout causes defects in skeletal development, and miR-92a⁻/⁻ mice are born at sub-Mendelian ratios, indicating roles in neural crest and mesoderm-derived lineages^85,86^. 92i in human 3D blastoids, which closely recapitulate the miRNA landscape of the E6 human blastocyst^6^, modestly reduced blastoid formation. Those blastoids that did form retained normal morphology, with embryonic and extra-embryonic lineages and a cavity at the end timepoint, suggesting that 92i may induce a developmental delay across the blastoid differentiation trajectory, echoing the lag seen in RACLs. Second, the embryo-level distribution of hsa-miR-92a-3p, which is abundant from the oocyte through the blastocyst stage and present in both TE and ICM as also previously reported^3,87^, underscores its relevance across early embryonic and extra-embryonic lineages. The specificity of the 92i phenotype is further supported by coherent de-repression of validated 3′UTR targets that, in human embryos, are predominantly highly expressed in epiblast or TE rather than hypoblast or EXM populations. Gene-set–level shifts in developmental and ECM programs in 92i RACLs reinforce this interpretation. We would like to note that RACL cultures are heterogeneous and contain cells spanning epiblast-intermediate, HLC, and EXMC states^22^, and bulk transcriptomic measurements may therefore obscure possible cell-type-specific responses to hsa-miR-92a-3p inhibition.

ESC miRNAs frequently act by tuning pluripotency signaling networks and cell-cycle machinery^14^. Consistent with this, we observed shifts in cell-cycle programs along the naïve hESC-RACL-NACL trajectory, and these programs were further altered under 92i conditions. At D7, 92i RACLs contained fewer dividing cells (in G2/M) and retained an earlier cell-cycle signature, with increased proportions in S compared to controls, in line with prior links between the miR-17∼92 cluster and proliferation and self-renewal^13,14^. Interestingly, both miR-17∼92 and the paralogous miR-106-363 cluster are up-regulated in paused ESCs^88^. Recent work implicates miR-92a in regulating dormancy during embryonic diapause^88^, a reversible state of suspended blastocyst development characterized by minimal cell proliferation and metabolism^89^. In mice, diapause occurs at the blastocyst stage when implantation is temporarily halted^90^, and miR-92a overexpression prolongs blastocyst survival in vitro^88^. Conversely, knockdown of miR-92a-3p in zebrafish ovaries increases developmental arrest at the 1-cell stage^91^, suggesting species-, context-, or miRNA arm usage-specific functions during early development. In mice, FGF/ERK signaling regulates primitive endoderm (mouse hypoblast equivalent) specification not only through directly altering enhancers or endoderm priming but also by coordinating inheritance of cell-cycle lengths^71,72,92^, highlighting a mechanistic interface between signaling pathways and proliferative timing. Among the up-regulated targets, *FGF2* emerged as one of the most highly expressed, derepressed genes following hsa-miR-92a-3p inhibition. Temporal RNA and protein analyses revealed a window during which FGF ligand availability and receptor composition influence lineage progression. The 92i phenotypes we observed, including reduced numbers of GATA6^+^/NANOG^-^, FGF2^+^/GATA6^+^, and GATA6^+^/FGFR1^+^ cells and altered FGF2 localization, were consistent with incomplete engagement of the HLC and EXMC trajectory. These results accord with recent work identifying FGF signaling as a central driver of hypoblast and HLC specification in human blastocysts as well as in 2D and 3D stem-cell–based models^62,63^. Collectively, our findings support a model in which hsa-miR-92a-3p coordinates cell-cycle phase transitions with fate decision, an established yet underexplored axis of early human cell lineage specification^93^. We propose that this coordination is mediated, at least in part, through FGF signaling, specifically through modulation of ligand levels, receptor switching, and matrix associated interactions, allowing hsa-miR-92a-3p to act as a timing regulator that links proliferative state to hypoblast and EXM fate acquisition.

## MATERIALS AND METHODS

### hESC lines and culture conditions

Naïve H9 (WA09, Wicell) hESCs (female), which were previously converted from primed hESCs using the NaïveCult Induction kit (STEMCELL Technologies), were cultured in NaïveCult Expansion Medium (STEMCELL Technologies) in 5% O2/5% CO2 at 37°C. Naive hESCs were dissociated with TrypLE Express (Thermo Fisher Scientific) every 2–5 days and re-plated on mitomycin C treated (Thermo Fisher Scientific) or irradiated (Merck) CF1 MEF feeders, prepared a day before hESC seeding. The cell culture medium was supplemented with 10 μM ROCKi Y-27632 (Selleckchem) for the first 24 h following plating. Primed H9 hESCs (WA09, Wicell) were cultured in Essential 8 medium (Thermo Fisher Scientific) on Geltrex (Thermo Fisher Scientific) coated tissue culture dishes in 5% CO2 at 37°C. The cells were passaged every 3–5 days after a 3-min incubation with 0.5 mM EDTA (Thermo Fisher Scientific). The potential influence of female genetic background on the observed phenotypes was not evaluated in this study. Cell lines obtained from Wicell were rigorously authenticated by the supplier and routinely confirmed to be mycoplasma-negative.

### Human embryos and ethical issues

Collection and experiments using human embryos were approved by the Helsinki University Hospital Ethics Committee (diary number HUS/1069/2016) and research permission was approved by the Helsinki University Hospital Research Committee. Couples who had undergone infertility treatment at the Helsinki University Hospital Reproductive Medicine unit were offered the option to donate their cryopreserved embryos for research after termination of infertility treatments and expiration of embryo preservation timespan. Surplus human embryos were donated with informed consent, and donors were explicitly informed that donating embryos to research was voluntary. Patients received information about the study and experimental procedures, were offered counselling, and were not compensated for embryo donation. No human embryos were created or manipulated for research purposes. Embryo culture was terminated at the blastocyst stage, that is, several days prior to the expected formation of the primitive streak or onset of gastrulation. The work was conducted in strict accordance with local legislation, ethical guidelines and regulations, and the principles outlined in the International Society for Stem Cell Research guidelines.

Slow-frozen day 2 embryos, donated for research purposes, were thawed using RapidWarm Omni (Vitrolife). Two different methods were used. In the first method 500 µl of the Omni solutions 2, 3 and 4 were equilibrated to room temperature (RT). Ampoules were thawed in a water bath (+37°C) for 90 seconds. The contents of the ampoules were transferred to a petri dish; the embryos were promptly moved into Omni 2 for 5 min, RT. Subsequently, the embryos were transferred to Omni 3 for 5 min, RT, and then the embryos were transferred to Omni 4 for 5 min, RT. After rinsing in culture medium (Gx-TL, Vitrolife), the embryos were cultured in GX-TL until day 6. In the second method (one-step method) 1 ml of Omni Warm 2 was warmed to 37°C in ambient atmosphere. Ampoules were thawed in a water bath (+37°C) for 90 seconds. The contents of the ampoules were transferred to a petri dish; the embryos were promptly moved into Omni 2. After 1 min (+37°C) the embryos were rinsed in culture medium (Gx-TL, Vitrolife). The embryos were cultured in GX-TL until day 6.

### RACL and NACL differentiation

RACL and NACL differentiations were done as described in Linneberg-Agerholm et al.^21^ Naïve hESCs were plated onto feeders at 5 x 10^4^ cells/cm² and cultured in RACL [RPMI 1640 medium with GlutaMAX (Thermo Fisher Scientific), B-27 minus insulin (Thermo Fisher Scientific), 10 ng/ml leukemia inhibitory factor (LIF; Qkine), 3 µM Laduviglusib (CHIR99021; Selleckchem), and 100 ng/ml Activin A (Qkine)] for 7–8 days. The medium was changed every other day until cells reached confluence, after which it was replaced daily. When RACLs reached confluence, cells were passaged by Accutase (Gibco) dissociation, detached from the plate using a cell scraper, and collected as clusters using a stripette. The cells were re-plated onto feeders in NACL (N2B27 supplemented with 100 ng/ml ActA, 3 µM Laduviglusib and 10 ng/ml LIF) every 3–5 days and supplemented with 10 µM ROCKi Y-27632 (Selleckchem) for the first 24 h. NACLs were frozen in 60% NACL supplemented with 10 µM ROCKi Y-27632 (Selleckchem), 30% foetal bovine serum (FBS; Thermo Fisher Scientific) and 10% dimethyl sulfoxide (DMSO).

### Blastoid differentiation

Blastoids were generated according to Heidari Khoei et al.^94^ Naïve H9 hESCs were dissociated with Accutase (Gibco), inactivated in 0.1% bovine serum albumin (BSA) (Biowest)–DMEM/F12 (Thermo Fisher Scientific), passed through a 40-µm cell strainer (Corning) and centrifuged at 300g for 5 min. Cells were plated onto 0.1% gelatin-coated dishes in NaïveCult Expansion Medium (STEMCELL Technologies) with 10 μM ROCKi Y-27632 (Selleckchem) and incubated in 5% O2/5% CO2 at 37°C for 75 min. Non-adherent cells were collected, resuspended in N2B27 basal medium and plated onto Aggrewell 400 (STEMCELL Technologies) plates at 6 × 10^4^ cells per well in 500 µl N2B27 with 10 µM ROCKi Y-27632, 0.3% BSA and 1× penicillin–streptomycin (BioNordika). Medium was changed 24 h after plating (day 1), and 500 µl of 2× PALLY medium [1 x PALLY: N2B27 supplemented with 1 µM PD0325901 (MedChemExpress), 1 µM A83-01 (MedChemExpress), 10 ng/ml LIF (Qkine), 3 µM lysophosphatidic acid (Tocris Bioscience), 10 µM ROCKi Y-27632 and 1× penicillin–streptomycin] was added to each well. Half of the medium was replaced with fresh 1×PALLY medium on day 2. On day 3 and 4, 500 µl of the medium was removed and 800 µl of LY medium [N2B27 supplemented with 500 nM LPA (Tocris Bioscience), 10 µM ROCKi Y-27632 (Selleckchem), and 1× penicillin–streptomycin] was added. Blastoid formation was completed on day 5.

### Inhibitor treatment

After 24 h of RACL differentiation, cells were treated with a hsa-miR-92a-3p targeting, a seed mismatch control or a non-targeting scramble control In vivo miRCURY LNA microRNA inhibitor (QIAGEN). RACL medium and the LNAs were changed every 24 h until cells were fixed or collected for analyses. During blastoid generation the LNA treatment was initiated upon plating naïve hESCs onto Aggrewell plates and the LNAs were changed every 24 h until cells were fixed or collected for analyses. Both 5’ fluorescein (FAM)-labelled and unlabeled LNAs were utilized in the experiments. The LNAs had the following sequences: NegA: ACGTCTATACGCCA; I-hsa-mir-92a-3p: CGGGACAAGTGCAAT; I-hsa-mir-92aMM: CGGAGCAAGTACGAT

### Immunocytochemistry and imaging

Naïve and primed H9 hESCs, RACLs, and NACLs were fixed on Ibidi four- or eight-well μ-slides with 4% paraformaldehyde in PBS at RT for 10–15 min. The cells were washed three times in DPBS (Thermo Fisher Scientific) and permeabilized in 0.5% Triton X-100 (Sigma Aldrich) in DPBS at RT for 7 min, followed by a single wash. Unspecific primary antibody binding was blocked using Blocking solution (AH Diagnostics) at RT for 10 min. Primary antibodies anti-Fibroblast Growth Factor-Basic (1:500; Sigma Aldrich), anti-FGF Receptor 1 (1:500; Cell Signaling Technology), anti-FGF Receptor 2 (1:800; Cell Signaling Technology), anti-Vimentin antibody (1:1000; Abcam), anti-GATA4 (1:400; Thermo Fisher Scientific), anti-GATA6 (1:500; R&D Systems), anti-GATA3 (1:200; Thermo Fisher Scientific), anti-NANOG (1:200; Cell Signaling Technology), anti-OCT3/4 (1:500, Santa-Cruz Biotechnology), anti-human SUSD2 clone W5C5 (1:500; BioLegend), anti-KLF17 (1:500; Sigma Aldrich) and anti-Cytokeratin 7 Monoclonal Antibody (1:300; Thermo Fisher Scientific) were diluted in 0.1% Tween-20 in DPBS and incubated overnight at 4 °C. The cells were washed three times in 0.1% Tween-20 in DPBS and incubated with secondary antibodies [anti-mouse Alexa Fluor 647 (Thermo Fisher Scientific, #A31571), anti-rabbit Alexa Fluor 647 (Thermo Fisher Scientific, #A32795), anti-rat Alexa Fluor 594 (Thermo Fisher Scientific, #A21209), anti-goat Alexa Fluor 594 (Thermo Fisher Scientific, #A11058), anti-goat Alexa Fluor 488 (Thermo Fisher Scientific, #A32814), anti-mouse Alexa Fluor 594 (Thermo Fisher Scientific, #A21203) and anti-rabbit Alexa Fluor 594 (Thermo Fisher Scientific, #A21207)] diluted 1:1000 in the same buffer at RT for 30 min. After two washes with 0.1% Tween-20 (Thermo Fisher Scientific), nuclei were counterstained with DAPI (Invitrogen; 1:1000 in DPBS) and Alexa Fluor 488 Phalloidin (Thermo Fisher Scientific; 1:1000 in DPBS) for 10 min, RT. Cells were imaged with Andor Dragonfly 505 confocal microscope, and the images were captured with Apo LWD 40x/1.15NA glycerol objective and Plan Apo VC 60x/1.2NA water immersion objective or with Leica TCS SP8 X white light laser confocal microscope and captured with HC PL APO CS2 40x/1.30NA oil objective or Leica SP8 CARS HC PL APO CS2 40x/1.10NA water objective. The images were processed using Fiji. A Gaussian blur (σ = 1.0) was applied in ImageJ to images presented in Figure 2 and Supplementary Figure 2. Nuclear segmentation was performed using Fiji on the DAPI channel following Gaussian blur filtering (σ = 2). Nuclei were segmented using Otsu thresholding to generate binary masks, and fluorescence intensities were quantified and exported for analysis. Positive and negative nuclei were defined using empirically determined intensity thresholds based on inspection of DAPI-normalized signal distributions and corresponding fluorescence images.

### Blastocyst and blastoid immunostaining and imaging

Blastocysts and blastoids were fixed with 4% PFA for 15 min, RT, washed three times with 0.1% Tween20-DPBS (PBST) and permeabilizated in 0.3% (blastoids) or 0.5% (blastocysts) Triton X-100 in DPBS for 30 min, RT, followed by a wash in PBST for 5–10 min. Unspecific antibody binding was blocked with 10% donkey serum in 0.3% Triton X-100 in DPBS for 2 h at RT (blastoids) or with commercial Blocking solution (AH Diagnostics) for 10 min, RT (blastocysts). Samples were incubated with primary antibodies diluted (GATA3 1:200, NANOG 1:200, GATA6 1:600, FGF2 1:500, FGFR1 1:500, FGFR2 1:800) in PBST overnight at +4C. After primary antibody incubation samples were washed three times in PBST at RT for 5-10 min, and treated with secondary antibodies (anti-goat Alexa Fluor 488; anti-rabbit Alexa Fluor 594; anti-rabbit Alexa Fluor 647; anti-mouse Alexa Fluor 594; anti-mouse Alexa Fluor 647) diluted 1:1000 (blastoids) or 1:500 (blastocysts) in PBST for 2 h in dark, RT. Samples were washed three times in PBST at RT for 5-10 min and counterstained with DAPI 1:1000 in PBST for 15 min, at RT. Blastocysts were imaged with Leica TCS SP8 X white light laser confocal microscope. The images were captured with HC PL APO CS2 40x/1.30NA oil objective using 1,024 x 1,024 scan format, 2µm z-step size and 1.5x zoom. The data were processed using Fiji^95^. Blastoids were imaged with Leica Thunder Imager 3D Cell Culture microscope. The images were captured with 40x air, HC PL Fluotar, NA 0.8. Lumencor Spectra X LED light source with wavelengths: 395nm, (filter 395/25), 470nm, (filter 475/28), 550nm, (filter 555/28), 640nm, (filter 640/30) with 1,024 × 1,024 scan format and 2× 2 binning. Images were acquired by taking a Z-stack with size of 1.00um using a Leica DFC9000GTC 4.2MPx sCMOS B&W camera and Leica LAS X 3.9.0 software. The data were processed using Fiji. The Z-stack was converted into a single-plane image using maximum intensity projection. The maximal widths of the ICM and the cavitation cavity were quantified from the resulting image to assess the ICM/cavity ratio. For the assessment of generation efficiency and size, blastoids were imaged with Leica Thunder Imager 3D Cell Culture system with a 10x, HC PL Fluotar, NA 0.32, air objective under brightfield settings. Images were captured with the Leica DFC9000GTC 4.2MP sCMOS B&W camera and Leica LAS X 3.9.0 software Navigator add-on. The Navigator add-on generates tile-scan images of the entire well, which are seamlessly stitched together with LAS-X software. Cavitated cell aggregates were measured for the maximum and minimum cavity diameter from which a mean was calculated. During the cavitation process, blastoids tend to float, causing them to displace from of the microwells. As a result, only the blastoids that remained at the bottom of the microwell were in focus for imaging and included in the final count. The data were processed using Fiji.

### Flow cytometry

Cells were detached using TrypLE (8 min at 37°C), centrifuged at 400 x g for 5 min, and resuspended in FACS buffer (5% FBS in DPBS). From this point on, cells were kept on ice. The cell pellet was fixed with 3.8% PFA diluted in FACS buffer for 10 min on ice. Cells were centrifuged at 400 x g for 5 min and the supernatant was removed. For cell-cycle profiling, one drop of FxCycle Violet Ready Flow Reagent (Thermo Fisher Scientific) was added to the PFA-fixed cells suspended in 500 μL FACS buffer and incubated for 30 min, RT. Novocyte Quanteon flow cytometer (Agilent) was used to measure DNA content. Cell-cycle analysis was performed using NovoExpress software (Agilent).

### RNA extraction and qPCR

Total RNA was isolated using the RNEasy mini kit (QIAGEN, #74104) according to the manufacturer’s protocol. For cDNA synthesis, 1μg of total RNA was reverse-transcribed by MMLV-RTase (Promega, #M1701) with oligo dT primers (Thermo Fisher), Random Hexamer Primer (Thermo Fisher), and Ribolock RNase inhibitor (Thermo Fisher). The reaction was incubated at 37°C for 90 min, then inactivated at 95°C for 5 min. The resulting cDNA was used as a template for qPCR using 5x HOT FIREPol qPCR Mix (Solis BioDyne) on the StepOnePlus System (Applied Biosystems/Thermo Fisher Scientific). Relative expression values were calculated using the 2^-ΔΔCt^ method, with GAPDH as an internal control. The used primer sequences are listed in Supplementary Table 15.

### sRNA extraction and miRNA qPCR

Blastoids were washed with DPBS and collected. Naïve hESC were washed, detached and plated onto 0.1% gelatin-coated dishes in NaïveCult Expansion Medium (STEMCELL Technologies) with 10 μM ROCKi Y-27632 (Selleckchem) and incubated in 5% O2/5% CO2 at 37 °C for 75 min. Non-adherent cells were collected. The samples were lysed in QIAZOL (QIAGEN), vortexed, and were further homogenized by passing the solution 5–10 times through a 21-gaugeneedle attached to a sterile plastic syringe. sRNA isolation was performed with the miRNeasy micro kit (QIAGEN) following the manufacturer’s instructions including the DNAse I treatment. sRNA was eluted in 20 µl of RNase-free water. An aliquot of 1 µl was taken for the quality control. sRNA integrity values were assessed by running 1 µl of sRNA solution in an RNA ScreenTape (Agilent Technologies) in the Agilent 4200 TapeStation System. qPCR validation of hsa-miR-92a-3p was performed using the miRCURY LNA™ miRNA PCR kit (QIAGEN, ref.339320) and Quantstudio 5 instrument (Applied Biosystems, Thermo Fisher Corporation). Three biological replicates from blastoids (n=3) and naïve hESCs (n=3) were analyzed along with a positive control sample and a negative sample (NTC). Human XpressRef Universal Total RNA (QIAGEN, ref.338112) was used as the positive control sample. 5ng/ul total RNA was used as input material. Spiked-in Uni-Sp6 was used as a technical control for cDNA synthesis. Assays for hsa-miR-92a-3p, hsa-miR-372-3p and hsa-miR-103-3p were analyzed using qPCR, with each sample run in triplicate for each assay. Relative expression values were calculated using the 2^-ΔΔCt^ method^96^, with hsa-miR-103-3p as an internal control.

### Bulk sRNA sequencing library preparation and sequencing

Naïve hESCs, RACLs and NACLs were collected by washing the cells once with DPBS and lysing them in QIAZOL (QIAGEN). RNA isolation was performed with the miRNeasy micro kit (QIAGEN) following the manufacturer’s instructions. RNA was eluted in 14 µl of RNase-free water. Library preparation from 10 ng of total RNA was performed according to QIAseq® miRNA Library Kit Handbook V. April 2021 (QIAGEN, Venlo, Netherlands). Library quality check was performed using Agilent Bioanalyzer High Sensitivity DNA assay (Agilent, Santa Clara, CA, USA) and Qubit HS DNA Assay (Thermo Fisher Scientific, Waltham, MA, USA) after which libraries were pooled for sequencing based on the results acquired from these assays. Libraries were sequenced on the Illumina NextSeq Mid Output 2×75bp flowcell (Illumina, San Diego, CA, USA).

### Bulk RNA sequencing library preparation and sequencing

Naïve hESCs, RACLs and NACLs, both LNA treated and untreated, were collected by washing the cells once with DPBS. RNA isolation was performed with the RNeasy mini kit (QIAGEN) following the manufacturer’s instructions including the DNAse I treatment. RNA was eluted in 30 µl of RNase-free water. Library preparation from 100 ng of total RNA was performed according to Illumina stranded total RNA prep with ribo-zero plus reference guide (Illumina, San Diego, CA, USA). Library quality check was performed using Agilent TapeStation 4200 (Agilent, Santa Clara, CA, USA) and Qubit HS DNA Assay (Thermo Fisher Scientific, Waltham, MA, USA) after which libraries were pooled for sequencing based on the results acquired from these assays. Sequencing was performed with Illumina NovaSeq6000 system using 2 lanes of S4 flow cell with lane divider (Illumina, San Diego, CA, USA). Read length for the paired-end run was 2×151 bp.

### Data and code availability

This paper analyzes publicly available data and previously unpublished data that is made available upon publishing. The Paloviita et al.^2^ human oocyte and embryo sRNA-seq data is available at FEGA Finland: EGAD50000000227. The Yang et al.^27^ human oocyte and embryo sRNA-seq data was acquired with accession number Gene Expression Omnibus (GEO): GSE95218. The Yang et al.^28^ mouse bulk sRNA-seq data of oocytes and embryos was retrieved with accession number: SRA: SRP045287. The sc-sRNA-seq data of human pre-implantation embryos^3^ was retrieved with accession number GEO: GSE249708. The scRNA-seq data of RACLs and NACLs^22^ was retrieved with accession number GEO: GSE204819. The RNA-seq and sRNA-seq data generated in this study are publicly available as of the date of publication. All original code is publicly available as of the date of publication.

### Pre-processing of sc/embryo sRNA-seq

Pre-processing and expression profiling of raw sRNA-seq reads from Paloviita et al.^2^ and Yang et al.^27^ were performed using sRNAbench^97,98^ following the approach used in Paloviita et al.^2^ Reads containing at least the first 10 nucleotides of the adapter sequence were adapter-trimmed and reads between 17–30 nucleotides in length were retained. Expression profiling was carried out using the genome-mode workflow of sRNAbench^97^. Briefly, pre-processed reads were aligned with Bowtie (v.1.3)^99^ to the human genome (GRCh38, primary assembly, Ensembl), allowing one mismatch. Reads were then sequentially mapped to the following reference databases: human miRBase v22^81^, Ensembl cDNA (hg38), Ensembl noncoding RNA (hg38), RNAcentral v14 (hg38), and os-piRNA^27^. A read was assigned to a reference RNA only if its alignment matched the reference sequence and its genomic coordinates fell entirely within those of the annotated RNA. The sum of single-assignment reads reported by sRNAbench was used for each miRNA to avoid inaccuracies introduced by multimapping reads. miRNAs were retained for downstream analysis only if they had detectable expression (read count > 0) in at least three samples. Read counts for all retained miRNA species were normalized using the TMM method implemented in the edgeR (v3.42.4) R package^100^, with effective library sizes calculated from the filtered miRNA count matrix.

### Pre-processing of bulk sRNA-seq data

The Yang et al.^28^ sRNA-seq reads were pre-processed with trimgalore (v.0.6.7) keeping adapter clipped reads of 17–32 bp that were aligned to the mouse genome (GRCm38) and mouse reference miRNAs (miRBase v22) using Bowtie with parameters -n 0 -l 19 --best --strata --norc -M 1 -S. Samtools (v.1.10)^101^ was used to sort, index and quantify (idxstats) the aligned reads. miRNAs were retained for downstream analysis only if they had detectable expression (read count >= 1) in at least two samples. Read counts for all retained miRNA species were normalized using the TMM method, with effective library sizes calculated from the filtered miRNA count matrix. The naïve hESC, RACL, and NACL sRNA-seq reads were pre-processed with umitools (v.1.1.2) and trimgalore keeping adapter clipped reads of 17–32 bp that were de-duplicated (--method=unique) after alignment to the human genome (GRCh38) and human reference miRNAs (miRBase v22) using Bowtie^99^ with parameters -n 0 -l 19 --best --strata --norc -M 1 -S. Samtools^101^ was used to sort, index and quantify (idxstats) the de-duplicated and aligned reads. miRNAs were retained for downstream analysis only if they had detectable expression (read count >= 10) in at least two samples. Read counts for all retained miRNA species were then normalized using the TMM method, with effective library sizes calculated from the filtered miRNA count matrix.

### Pre-processing of bulk RNA-seq data

The bulk RNA-seq reads of naïve hESC, D4/D7-RACLs (medium, NegA, 92MM, 92i), and NACLs were pre-processed with trimgalore using parameter –paired and aligned to the human reference genome (GRCh38) with GENCODE (v41) transcript annotations and quantified using STAR (v.2.7.11a) with parameters --quantMode GeneCounts --outFilterType BySJout --outFilterMultimapNmax 20 --alignSJoverhangMin 8 --alignSJDBoverhangMin 1 --outFilterMismatchNmax 999 --outFilterMismatchNoverReadLmax 0.04 --alignIntronMin 20 --alignIntronMax 1000000 --alignMatesGapMax 1000000. Genes were retained for downstream analysis only if they had detectable expression (read count >= 10) in at least two samples. Read counts for all retained genes were then normalized using the TMM method, with effective library sizes calculated from the filtered count matrix.

### sc-sRNA-seq quality control, integration and clustering

The miRNA reads in the sRNA expression matrix for Russell et al.^3^ were used for downstream processing with the R package Seurat (4.3.0). Cells with less than one miRNA feature and miRNA features with expression in less than three cells were removed. The data was filtered as in Russell et al.^3^ Briefly, cells were required to have at least 0.5 million sequenced reads, less than 25% of mitochondrial UMIs, and more than 100 detected miRNA molecules. The data was log-normalized and scaled batch-wise using SCTransform. The data was integrated using functions SelectIntegrationFeatures with 3000 features, PrepSCTIntegration, FindIntegrationAnchors and IntegrateData for method SCT. Uniform Manifold Approximation and Projection (UMAP) was implemented using top 15 PCs and cells were clustered using the Louvain algorithm and a resolution of 0.6. The code used for the analyses and visualizations are provided on Zenodo after publication.

### scRNA-seq quality control, clustering, and pseudotime

The filtered STARsolo output from the D7-RACL and NACL scRNA-seq dataset by Pham et al.^22^ was used for downstream analysis with Seurat. Cells expressing fewer than 200 RNA features and RNA features expressed in fewer than three cells were excluded. The data were log-normalized and scaled using SCTransform, regressing out mitochondrial read content (percent.mt) and cell-cycle effects estimated using the CellCycleScoring function. UMAP was implemented using the top 20 PCs, and cells were clustered using the Louvain algorithm with a resolution of 0.6. Single-cell pseudotime trajectories were inferred using the slingshot (v1.8.0) R package, which enables the reconstruction of lineage relationships in a low-dimensional embedding^65^. Pre-computed cell embeddings and cluster assignments from the Seurat analysis were used as input to the slingshot function, with the start cluster set to RACL.intermediate.1. Pseudotime values for individual cells were computed using the slingPseudotime function with na.rm = TRUE. Mean pseudotime across the two inferred trajectories was calculated using the rowMeans function. Code used for all analyses and visualizations will be made available on Zenodo upon publication.

### Differential expression analysis

DE analysis was performed using the DESeq2 (v.1.40.1) R package^36^ with the filtered count matrices. For the bulk sRNA-seq differentiation experiment, the naïve hESC, RACL, and NACL groups were compared pairwise. miRNAs with an FDR < 0.05 and an absolute log₂ fold-change > 1 were classified as significantly differentially expressed. For the naïve hESC, D4- and D7-RACL, and NACL bulk RNA-seq data, DESeq2 models were fitted while regressing out differentiation batch effects. Results from all comparisons against the naïve hESC reference were extracted and combined, after which adjusted P-values were recalculated across the merged results using the Benjamini–Hochberg method^102^. Genes with FDR < 0.05 and absolute log₂ fold-change > 1 were considered significantly differentially expressed. In the 92i experiment, D4- and D7-RACL bulk RNA-seq samples were grouped into treatment (92i at 0.5 µM and 0.75 µM) and control pools (medium, NegA at 0.1/0.5/0.75 µM, and 92i at 0.1 µM), based on PCA of all detected genes. The 92MM samples were excluded from the comparisons. The DESeq2 design formula included differentiation batch, timepoint, group (control vs. treatment), and the interaction between timepoint and group. DE results for D4 and D7 were extracted separately, and adjusted P-values were recalculated jointly across both timepoints using the Benjamini–Hochberg method. Features with FDR < 0.05 and absolute log₂ fold-change > 0.5 were considered differentially expressed.

### Gene set enrichment analysis

GSEA analysis of the Gene Ontology (GO) (2023) database ‘Biological Process’ class was performed using the clusterProfiler (v.4.8.1) R package^103^ gseGO function with pvalueCutoff = 0.05. For visualization, enriched GO terms were filtered based on GO hierarchy using a custom script. The clusterProfiler simplify function was applied with cutoff = 0.7, by = “first_GO_level”, and select_fun = min to reduce redundancy among enriched terms. The msigdbr (v. 7.5.1) R package^104^ was used to obtain the C3 (regulatory target gene sets) collection from the Molecular Signatures Database (MSigDB). Gene set enrichment was performed using the GSEA function from clusterProfiler. Analyses were carried out for DE genes in D4-RACL, D7-RACL, and NACL samples compared pairwise against naïve hESCs, as well as for the 92i experiment by comparing treatment and control groups separately for D4- and D7-RACLs (see methods DE section).

### Statistical tests and visualization

The following R packages were used to create visualizations of the results: RColorBrewer (v1.1-3), gplots (v.3.1.3), plotrix (v.3.8-2), scales (v.1.2.1), corrplot (v.0.92), viridis (v.0.6.2), ComplexHeatmap (v.2.16.0), circlize (v.0.4.15). UpSetR (v.1.4.0), enrichplot (v.1.20.0), pheatmap (v.1.0.12). The following R packages were used for general data analysis: data.table (v.1.14.8), tidyverse (v.2.0.0), SingleCellExperiment (v.1.22.0), scater (v.1.28.0), tradeSeq (v.1.14.0), and fields (v.14.1). The following R packages were used for genomic data analysis: biomaRt (v.2.56.1), GenomicRanges (v.1.52.0), org.Hs.eg.db (v.3.17.0). Correlation analysis was performed on log-transformed TMM-normalized counts (pseudocount = 1), using Spearman’s rank correlation with complete observations. PCA was performed using the FactoMineR (v.2.8) and factoextra (v.1.0.7) R packages on log-transformed, TMM-normalized counts (pseudocount = 1), using the *PCA* function with scale.unit = TRUE. For analyses involving differentially expressed genes, the filtered count matrix was normalized using the limma (v.3.56.2) R package voom function^105^ to account for differentiation-related batch effects, and the resulting voom-transformed values were used as input to the PCA function with scale.unit = FALSE. PCA of human embryo scRNA-seq data^56^, restricted to the cells reannotated by Okubo et al.^23^, together with our naïve hESC, D4/D7-RACL, and NACL bulk RNA-seq samples was performed as follows. Embryo data were TMM-normalized using edgeR to mirror bulk RNA-seq preprocessing. Both datasets were then filtered to retain only genes with TMM > 2 in all samples and present in both datasets. The embryo expression matrix was log-transformed and scaled, and PCA was computed using the PCA function. Our bulk RNA-seq samples were similarly log-transformed and scaled and subsequently projected onto the embryo-derived principal component space using the predict function. Fuzzy clustering was performed using Mfuzz (v.2.60.0) with 4 centers and a fuzzification parameter of 3.8. Hierarchically clustered (Euclidean distance, average linkage) heatmaps were generated with scaled TMM-normalized data with the overall expression level shown as the log-transformation of the mean expression level across samples with a pseudocount of 0.0001. Blastoid generation efficiency was assessed by quantifying blastoids that exhibited a visible cavity and reached a mean diameter of at least 150 µm^68^. To compare the ICM–to-cavity ratio and blastoid generation efficiency between 92i-treated and control (NegA) conditions, pairwise Wilcoxon rank-sum tests were performed using the compare_means function in R. Statistical significance of differences in the numbers of positive and negative nuclei staining between the treatment and control groups was assessed by comparing DAPI-normalized fluorescence signal distributions. Pairwise Wilcoxon rank-sum tests were performed using the compare_means function in R. Percentages of cells in each cell-cycle phase, as determined by cell-cycle analysis using NovoExpress software (Agilent), were compared between treatment and control groups using pairwise Wilcoxon rank-sum tests implemented with the compare_means function in R. For analysis of qPCR data Kruskal-Wallis test with post-hoc Dunn’s test were calculated using GraphPad Prism 10.6.1. P-values < 0.05 were considered statistically significant. Information on replicate numbers and definition of center, and dispersion measures are indicated in the figure legends. The code used for the analyses and visualizations are provided on Zenodo after publication.

## Acknowledgements

We are grateful to the couples that have donated their surplus embryos for this project. We thank the staff at the Reproductive Medicine Unit of the Helsinki University Hospital for recruiting the couples to the oocyte and embryo donation program, Prof. Vincent Pasque for qPCR primers, and Dr. Madeleine Linneberg-Agerholm and Prof. Joshua Brickman for help with the RACL and NACL differentiation protocols. We acknowledge Biomedicum Imaging, Biomedicum Flow Cytometry, Biomedicum Functional Genomics, Institute for Molecular Medicine Finland (FIMM) Genomics NGS Sequencing, and HiPREP Core at FIMM Technology Centre units, University of Helsinki, with the support of the Helsinki Institute of Life Science (HiLIFE) and Biocenter Finland and the CSC–IT Center for Science Ltd. This study was supported by grants from the Research Council of Finland (Academy Fellowship grants #348111 and #353549), Sigrid Jusélius Foundation, Jane and Aatos Erkko Foundation, and Helsinki Institute of Life Science (HiLIFE fellowship), all to S.V. P.P. is supported by the Finnish Cultural Foundation, Orion Research Foundation sr, Finnish Fertility Society, and the Biomedicum Helsinki Foundation. S.N. is supported by University of Helsinki Doctoral Program in Biomedicine, the Finnish Fertility Society, and Instrumentarium Science Foundation.

## Author contributions

P.P. and S.V. conceived the study; S.V. supervised the project; P.P., E.H., R.G., I.K., S.N., R.P., and R.S. performed cell culturing; E.H., R.G., I.K., S.N., R.P., and R.S. performed immunostainings and confocal microscopy; E.H. and H.G. conducted flow cytometry analyses; P.P. and R.G. performed RNA and sRNA extractions; E.H. and R.G. performed qPCR; R.P. and R.S. conducted blastoid experiments; C.H.-G. and T.T. collected and thawed the embryo samples; P.P. performed bioinformatics analyses; P.P., E.H., and S.V. wrote the manuscript; and all authors edited and accepted the manuscript.

---

**Extended Data Figure 1.**
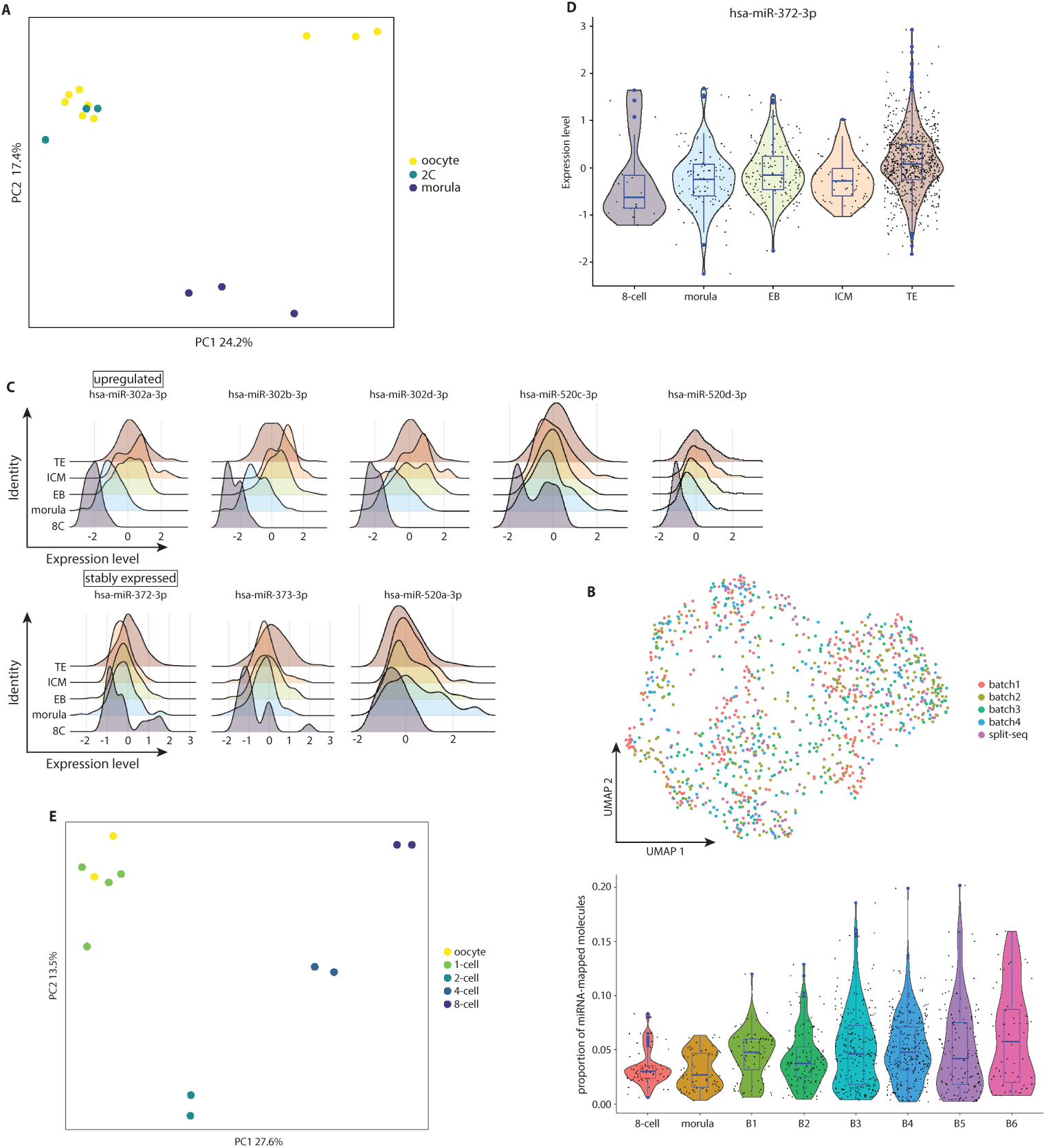
**(a)** PCA of log-transformed, TMM-normalized expression of all detected miRNAs in human oocytes and early embryos^27^, colored by developmental stage. **(b)** UMAP of Russell et al.^3^ data after batch correction with single cell small-RNA-seq batches and the split-seq sample indicated by color (top) and the proportion of miRNA-mapped molecules per morphological embryo grade shown as a blue boxplot overlaid on a violin plot (bottom). Boxplots show the median, 25^th^ and 75^th^ percentiles, and outliers; black dots indicate individual cells. **(c)** Ridge plot of miRNA molecule–proportion distributions (over total miRNA molecules) for fam-372 members in the Russell et al.^3^ dataset, grouped as up-regulated, down-regulated, or stable between 8-cell stage and inner cell mass or trophectoderm. See Figure 1e. **(d)** Boxplot (blue) overlaid on a violin plot showing normalized hsa-miR-372-3p expression across human embryo stages and lineages^3^, colored by group. Boxplots show the median, 25^th^ and 75^th^ percentiles, and outliers; black dots indicate individual cells. **(e)** PCA of log-transformed, TMM-normalized expression of all detected miRNAs in mouse oocytes and early embryos^28^, colored by developmental stage. **Abbreviations:**(1C) one-cell stage embryo; (2C) two-cell stage embryo; (4C) four-cell stage embryo; (8C) eight-cell stage embryo; (EB) embryoid body; (ICM) inner cell mass; (TE) trophectoderm; (B) blastocyst stage; (fam-372) seed family miR-302-3p/372-3p/373-3p/520-3p.

**Extended Data Figure 2.**
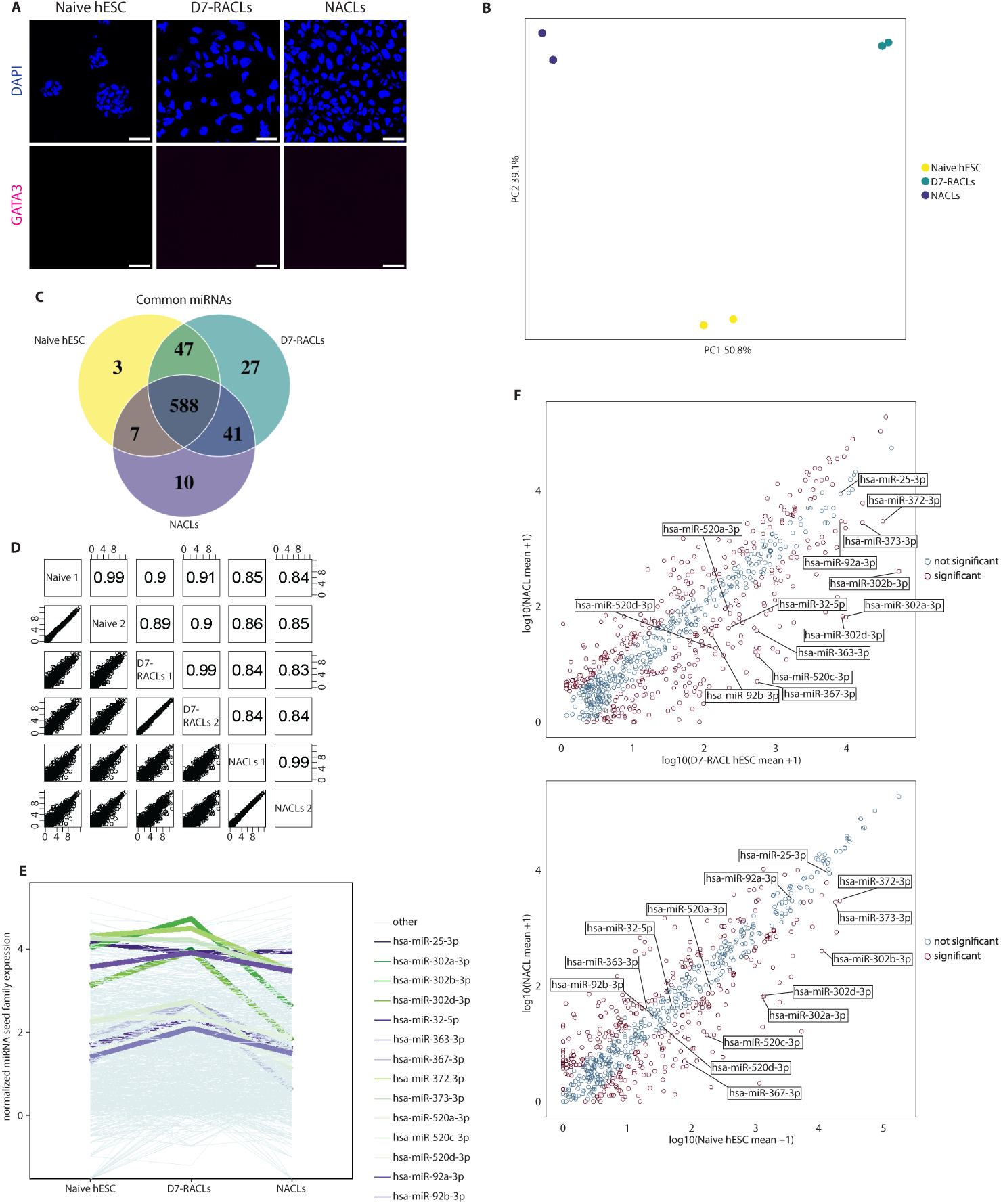
**(a)** Naïve H9 hESCs, D7-RACLs and NACLs immunostained with antibody recognizing GATA3 (red, not detected), and DNA counterstained with 4′,6-diamidino-2-phenylindole (DAPI). Scale bars 50 μm. **(b)** PCA of log-transformed, TMM-normalized expression of all detected miRNAs in naïve H9 hESCs, D7-RACLs, and NACLs (n = 2), colored by sample type. See Figure 2c. **(c)** Venn diagram of detected miRNAs in naïve H9 hESCs, D7-RACLs, and NACLs. **(d)** Pairwise Spearman correlations of log-transformed, TMM-normalized miRNA expression in naïve H9 hESCs, D7-RACLs, and NACLs, using only miRNAs detected in both samples. Scatter plots show expression values. **(e)** Log-transformed, mean TMM-normalized expression of fam-92 and fam-372 miRNAs in naïve H9 hESCs, D7-RACLs, and NACLs, colored by miRNAs of interest (others in grey). **(f)** Scatter plots of log-transformed, mean TMM-normalized miRNA expression comparing NACLs and D7-RACLs (n = 2; top) and naïve H9 hESCs and NACLs (n = 2; bottom). Significantly differentially expressed miRNAs (DESeq2: |log₂FC| > 1, FDR < 0.05) are shown in red; non-significant miRNAs in grey. Members of fam-92 and fam-372 are labeled. **Abbreviations:** (D7-RACLs) day 7 RACL cells; (NACLs) NACL cells; (fam-92) seed family miR-25-3p/32-5p/92-3p/363-3p/367-3p; (fam-372) seed family miR-302-3p/372-3p/373-3p/520-3p.

**Extended Data Figure 3.**
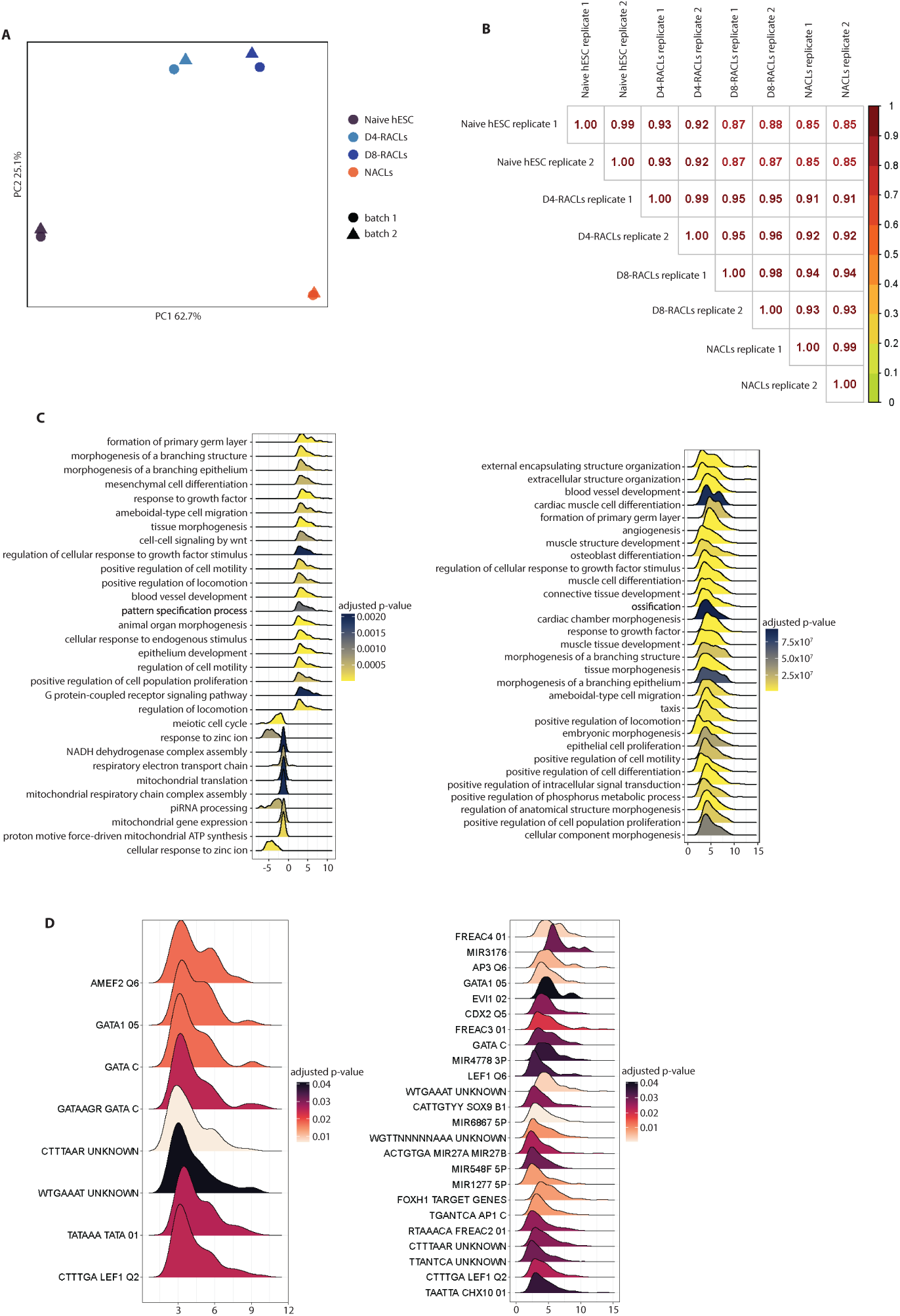
**(a)** PCA of log-transformed, TMM-normalized expression of differentially expressed genes (DESeq2: |log₂FC| > 1, FDR < 0.05) in naïve H9 hESCs, D4- and D7-RACLs, and NACLs (n = 2). Dot color indicates sample type; dot shape indicates differentiation batch. See Figure 2h. **(b)** Pairwise Spearman correlations of log-transformed TMM-normalized gene expression in naïve H9 hESCs, D4-and D7-RACLs, and NACLs, using only genes detected in both samples. **(c)** Enriched biological processes among differentially expressed genes for D4-RACLs vs. naïve H9 hESCs (left) and NACLs vs. naïve H9 hESCs (right), filtered for Gene Ontology term hierarchy. Color indicates adjusted p-value; x-axis shows log-fold-change distribution. **(d)** Enriched transcription factor and miRNA regulatory target gene sets among differentially expressed genes for D4-RACLs vs. naïve H9 hESCs (left) and D7-RACLs vs. naïve H9 hESCs (right). Color indicates adjusted p-value; x-axis shows log-fold-change distribution. **Abbreviations:** (D4-RACLs) day 4 RACL cells; (D7-RACLs) day 7 RACL cells; (NACLs) NACL cells.

**Extended Data Figure 4.**
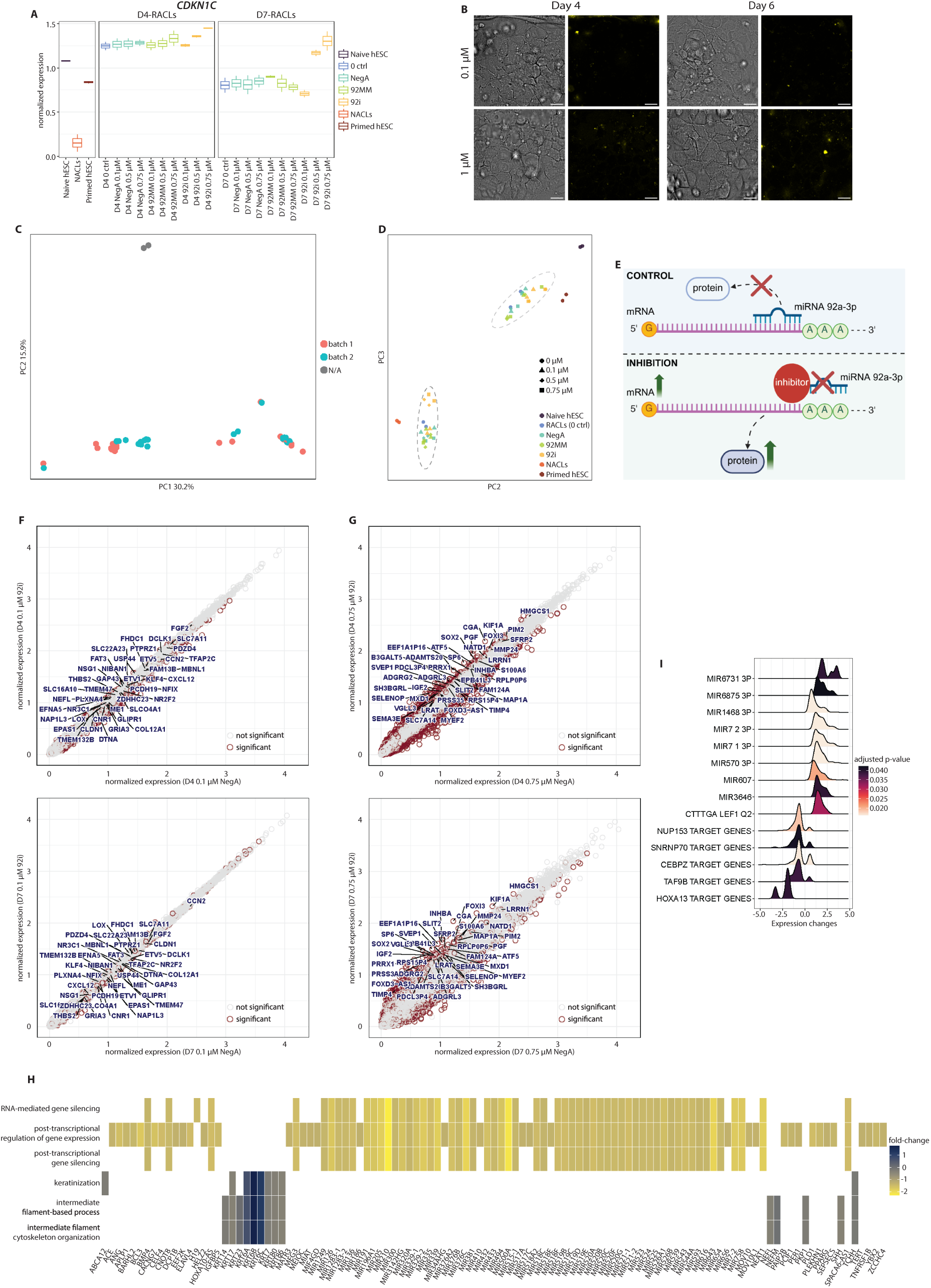
**(a)** Log-transformed, mean TMM-normalized *CDKN1C* expression measured by RNA-seq in D4- and D7-RACLs that were untreated (0 ctrl) or treated with 92i, 92MM, or NegA at the indicated doses, as well as in primed and naïve H9 hESCs and NACLs (n = 2). Colors indicate cell type or RACL treatment group. Boxplots show the median, the 25^th^ and 75^th^ percentiles. **(b)** Confocal images of D4-RACLs (left) and D6-RACLs (right) treated with a non-targeting FAM-labelled LNA at 0.1 μM (top) or 1 μM (bottom). Each panel shows a representative bright-field image and a fluorescence image detecting FAM signal (yellow). Scale bars 20 μm. **(c)** Differentiation-batch annotations corresponding to the PCA shown in Figure 3c. **(d)** PCA (PC2 and PC3), as in Figure 3d, showing only bulk RNA-seq samples from this study: primed and naïve H9 hESCs, NACLs, D4- and D7-RACLs that were untreated (0 ctrl) or treated with 92i, 92MM, or NegA. Shapes indicate treatment dose; colors indicate cell type or treatment group for the RACL samples. Dotted ovals mark D4- (light grey) and D7-RACLs (dark grey). **(e)** Schematic illustrating expected effects of hsa-miR-92a-3p inhibition on transcript and protein levels of its targets, which under control conditions (medium or NegA) are post-transcriptionally downregulated via hsa-miR-92a-3p binding. **(f)** Scatter plots of log-transformed, mean TMM-normalized gene expression in D4-RACLs (n = 2; top) and D7-RACLs (n = 2; bottom) treated with 0.1 μM 92i or NegA. Highly expressed (mean expression in the considered samples > 600) and significantly up-regulated predicted hsa-miR-92a-3p targets (TarBase)^106^ are labeled. Significantly differentially expressed genes (see methods for sample groups; DESeq2: |log₂FC| > 0.5, FDR < 0.05) are shown in red; non-significant genes in grey. **(g)** Scatter plots of log-transformed, mean TMM-normalized gene expression in D4-RACLs (n = 2; top) and D7-RACLs (n = 2; bottom) treated with 0.75 μM 92i or NegA. Highly expressed (mean expression in the considered samples > 600) and significantly up-regulated transcripts that are not hsa-miR-92a-3p targets (according to TarBase) are labeled. Significantly differentially expressed genes (see methods for sample groups; DESeq2: |log₂FC| > 0.5, FDR < 0.05) are shown in red; non-significant genes in grey. **(h)** Differentially expressed genes between 92i and control D7-RACLs included in the enriched biological processes shown in Figure 3j. Color key indicates fold-change values. **(i)** Enriched transcription factor and miRNA regulatory target gene sets among differentially expressed genes for 92i and control D7-RACLs. Color indicates adjusted p-value; x-axis shows log-fold-change distribution. **Abbreviations:** (D4-RACLs) day 4 RACL cells; (D7-RACLs) day 7 RACL cells; (NACLs) NACL cells; (92i) hsa-miR-92a-3p inhibitor; (92MM) seed-mismatch control; (NegA) non-targeting control.

**Extended Data Figure 5.**
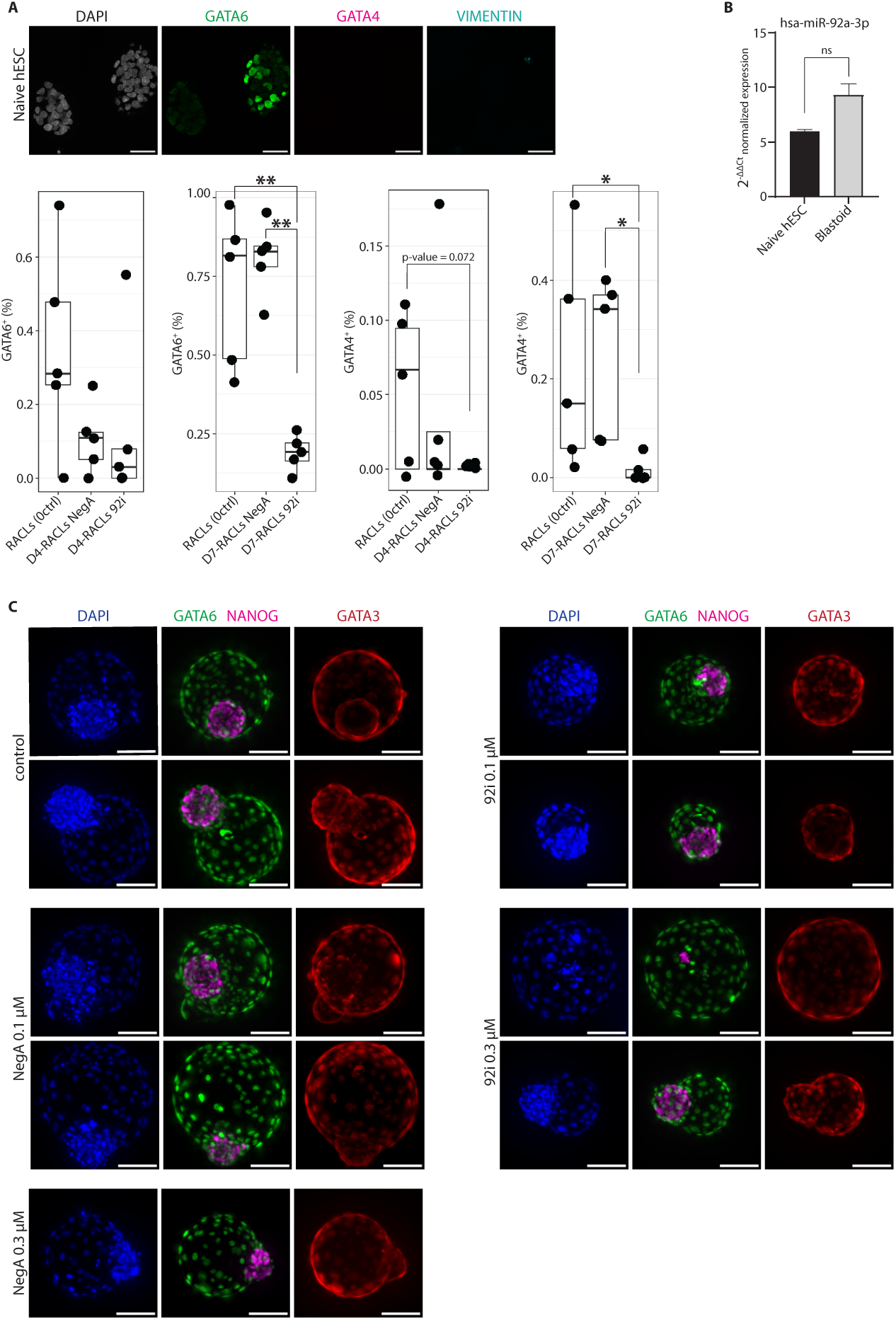
**(a)** Naïve H9 hESCs immunostained with antibodies recognizing FGF2 (green), GATA6 (magenta, not detected), NANOG (blue), and DNA counterstained with 4′,6-diamidino-2-phenylindole (DAPI). Scale bars 40 μm. **(b)** Naïve H9 hESCs, and D4- and D7-RACLs treated with 92i or NegA (0.75 μM) immunostained with antibodies recognizing FGFR1 (green), GATA6 (magenta), SUSD2 (blue), and DNA counterstained with 4′,6-diamidino-2-phenylindole (DAPI). Scale bars 40 μm. **(c)** Naïve H9 hESCs, and D4- and D7-RACLs treated with 92i or NegA (0.75 μM) immunostained with antibodies recognizing FGFR2 (green), GATA6 (magenta), SUSD2 (blue), and DNA counterstained with 4′,6-diamidino-2-phenylindole (DAPI). Scale bars 40 μm. **(d)** The percentage of GATA6-positive (⁺)/NANOG-negative (⁻) cells among all cells in D4- and D7-RACLs that were untreated or treated with 92i or NegA (0.75 μM). Quantification was performed from the images of the experiment for which representative images are shown in Figure 4d. Statistical significance was assessed using pairwise Wilcoxon rank-sum tests, with trends toward significance indicated in the plot. All other comparisons were not statistically significant. **Abbreviations:** (D4-RACLs) day 4 RACL cells; (D7-RACLs) day 7 RACL cells; (NACLs) NACL cells; (92i) hsa-miR-92a-3p inhibitor; (NegA) non-targeting control.

**Extended Data Figure 6.**
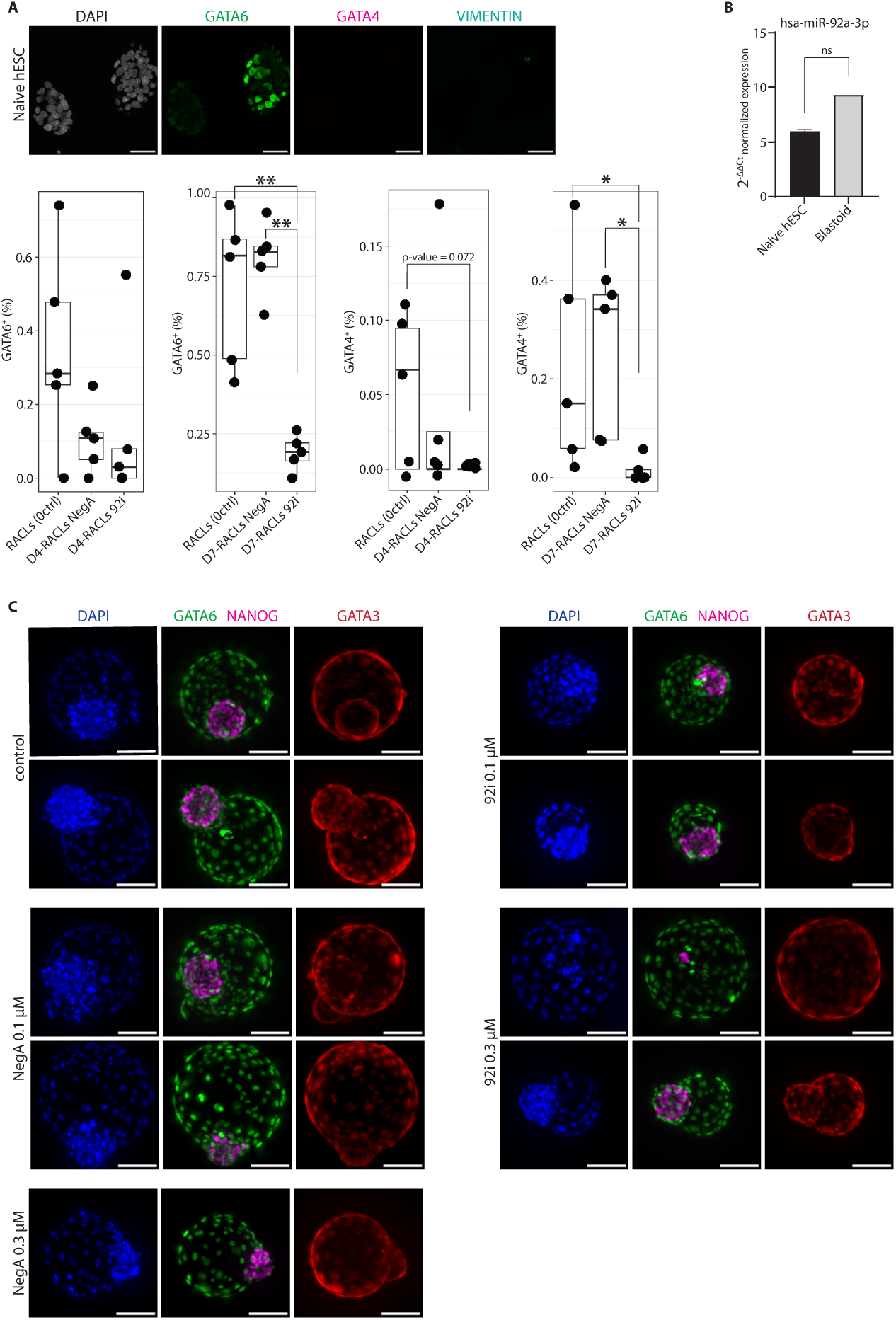
**(a)** Naïve H9 hESCs immunostained with antibodies recognizing GATA6 (green), GATA4 (magenta), vimentin (cyan), and DNA counterstained with 4′,6-diamidino-2-phenylindole (DAPI) (top). Scale bars 40 μm. The percentage of GATA6 positive (⁺)/ and GATA4⁺ among all cells in D4- and D7-RACLs that were untreated (0 ctrl) or treated with NegA or 92i (0.75 μM; bottom). Quantification was performed from the images of the experiment for which representative images are shown in Figure 5c. Statistical significance was determined using pairwise Wilcoxon rank-sum tests, with significant p-values indicated in the plot (*p < 0.05, **p < 0.01). **(b)** qPCR quantification of hsa-miR-92a-3p levels in naïve H9 hESCs and blastoids derived from naïve H9 hESCs. The barplot shows mean ± standard deviation (n= 3). Statistical significance was determined using Wilcoxon rank-sum test (p = 0.1, not significant (ns)). **(c)** Blastoids from naïve H9 hESCs that were untreated (control) or treated with NegA or 92i (0.1 and 0.3 μM) during differentiation immunostained with antibodies recognizing GATA6 (green), NANOG (magenta), GATA3 (red), and DNA counterstained with 4′,6-diamidino-2-phenylindole (DAPI; n=2). Scale bars 50 μm. **Abbreviations:** (D4-RACLs) day 4 RACL cells; (D7-RACLs) day 7 RACL cells; (NACLs) NACL cells; (92i) hsa-miR-92a-3p inhibitor; (NegA) non-targeting control.

**Extended Data Figure 7.**
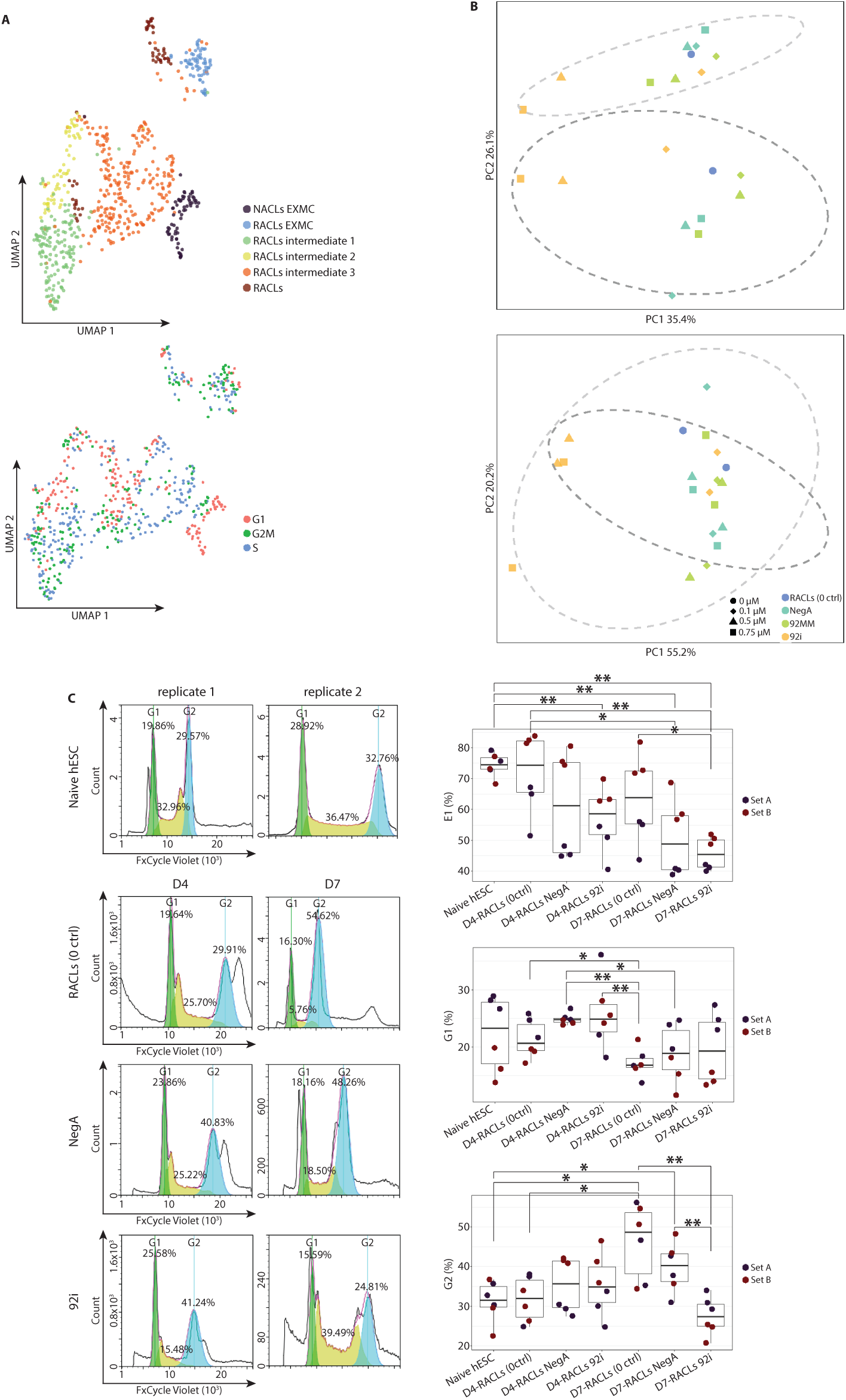
**(a)** UMAP of Pham et al.^22^ single cell RNA-seq data showing cell-type annotations (top) and Seurat-based cell-cycle phase assignments (bottom) for D7-RACLs and NACLs. **(b)** PCA of log-transformed, TMM normalized expression of Seurat’s cell-cycle genes in primed and naïve H9 hESCs, and NACLs, and D4-RACLs (top) and D7-RACLs (bottom) that were untreated (0 ctrl) or treated with 92i, 92MM, or NegA (n=2). Shapes indicate treatment dose; colors indicate cell type or treatment group for the RACL samples. Dotted ovals highlight samples from the two differentiation batches: batch 1 (light grey) and batch 2 (dark grey) **(c)** Cell-cycle profiles in naïve H9 hESCs and D4- and D7-RACLs that were untreated or treated with NegA or 92i (0.75 μM; left). DNA content was measured using FxCycle Violet and flow cytometry. Representative data are shown from two differentiation experiments (see Figure 6d for biological replicate) with three technical replicates per sample type (day and treatment). Percentage of cells included in the cell-cycle analysis (E1) and percentage of cells in the G1 and G2 phases for naïve H9 hESCs and RACL subgroups (right). The boxplot indicates the median, the 25^th^ and 75^th^ percentiles; dots represent replicates, colored by differentiation batch. Statistical significance was determined using pairwise Wilcoxon rank-sum tests, with significant p-values indicated in the plot (*p < 0.05, **p < 0.01). **Abbreviations:** (D4-RACLs) day 4 RACL cells; (D7-RACLs) day 7 RACL cells; (NACLs) NACL cells; (92i) hsa-miR-92a-3p inhibitor; (92MM) seed-mismatch control; (NegA) non-targeting control.

